# Decoding gene regulation in the mouse embryo using single-cell multi-omics

**DOI:** 10.1101/2022.06.15.496239

**Authors:** Ricard Argelaguet, Tim Lohoff, Jingyu Gavin Li, Asif Nakhuda, Deborah Drage, Felix Krueger, Lars Velten, Stephen J. Clark, Wolf Reik

## Abstract

Following gastrulation, the three primary germ layers develop into the major organs in a process known as organogenesis. Single-cell RNA sequencing has enabled the profiling of the gene expression dynamics of these cell fate decisions, yet a comprehensive map of the interplay between transcription factors and cis-regulatory elements is lacking, as are the underlying gene regulatory networks. Here we generate a multi-omics atlas of mouse early organogenesis by simultaneously profiling gene expression and chromatin accessibility from tens of thousands of single cells. We develop a computational method to leverage the multimodal readouts to predict transcription factor binding events in cis-regulatory elements, which we then use to infer gene regulatory networks that underpin lineage commitment events. Finally, we show that these models can be used to generate *in silico* predictions of the effect of transcription factor perturbations. We validate this experimentally by showing that Brachyury is essential for the differentiation of neuromesodermal progenitors to somitic mesoderm fate by priming cis-regulatory elements. The data set can be interactively explored at https://www.bioinformatics.babraham.ac.uk/shiny/shiny_multiome_organogenesis/

## Introduction

In mammals, specification of the basic body plan occurs during gastrulation, when the pluripotent epiblast is patterned to give rise to the three primary germ layers. Subsequently, these progenitors generate all major organ systems in a process known as organogenesis (Arnold and Robertson, 2009; Bardot and Hadjantonakis, 2020; Tam and Loebel, 2007). In the mouse, germ layer formation and early organogenesis have been profiled using a variety of genomics technologies, including single-cell RNA-sequencing (scRNA-seq), which led to the annotation of multiple cell types and the characterisation of differentiation trajectories (Cao et al., 2019; Ibarra-Soria et al., 2018; Pijuan-Sala et al., 2019). Some efforts to profile the epigenome during these stages have produced bulk chromatin accessibility using ATAC-seq and histone profiling with ChIP-seq at E7.5 (Xiang et al., 2020), single-nucleus (sn) chromatin accessibility maps at E8.25 with snATAC-seq (Pijuan-Sala et al., 2020) and single-cell transcriptome, nucleosome positioning and DNA methylation up to E7.5 with scNMT-seq (Argelaguet et al., 2019). These data demonstrate the dynamic remodelling that the epigenome undergoes during development. However, a comprehensive characterisation of the epigenome changes and the cis-regulatory elements involved in the transition from gastrulation to early organogenesis is still lacking, as well as an integration of this information with the transcriptome. Furthermore, the genomic positions and the target genes of the various transcription factors (TFs) that control these developmental trajectories have only been explored for a limited set of TFs and using *in vitro* systems. A catalogue of TF binding sites during mouse early organogenesis *in vivo* is lacking.

Single-cell multimodal technologies have huge potential for the study of gene regulation (Chen et al., 2019; Clark et al., 2018; Luo et al., 2022; Ma et al., 2020; Zhu et al., 2019, 2021). In particular, the ability to link epigenomic with transcriptomic changes allows the inference of gene regulatory networks (GRNs)(Aibar et al., 2017; Davidson and Erwin, 2006; Kamimoto et al., 2020; Kartha et al., 2021; Materna and Davidson, 2007). GRNs are able to capture the interplay between TFs, cis-regulatory DNA sequences and the expression of target genes (Garcia-Alonso et al., 2019; Levine and Davidson, 2005; Stadhouders et al., 2018), and can hold predictive power of cell fate transitions and gene perturbations (Kamimoto et al., 2020). Methods that derive GRNs from single-cell genomics data have been developed (Aibar et al., 2017; Fleck et al., 2021; Kamimoto et al., 2020; Kartha et al., 2021) and applied to the developing fly brain (Janssens et al., 2022) but similar analyses of mammalian development are lacking. In addition, GRN inference relies on accurate TF binding data, yet limited knowledge of TF binding exists for early embryonic development due to limitations in experimental methods such as ChIP-seq or CUT&RUN, which require large numbers of cells (Skene and Henikoff, 2017) and faithful antibodies. It is thus unrealistic to profile a large fraction of all TFs even in a single biological context (Lambert et al., 2018; Park, 2009). Instead, TF binding sites are typically inferred from the presence of a sequence motif within accessible chromatin (Castro-Mondragon et al., 2021; Schep et al., 2017; Weirauch et al., 2014). This approach can be successful for some TFs that display non-redundant DNA motifs with high sequence specificity, but the presence of a TF motif does not guarantee the existence of an active binding site (Wang et al., 2012). Moreover, the use of DNA motifs as a proxy for TF binding is not well suited for the study of TFs that share similar DNA motifs, and also for TFs linked to short motifs. Thus, alternative methods for predicting TF binding sites are required.

Recent technological advances have enabled the simultaneous profiling of RNA expression and epigenetic modalities from single cells at high-throughput (Chen et al., 2019; Ma et al., 2020; Zhu et al., 2019). This provides a unique opportunity to systematically decode the TF activities and the GRN structure that underpins cell fate transitions. Here, we perform snATAC-seq and snRNA-seq from the same nuclei from a time course of mouse embryonic development from E7.5 to E8.75. We develop a computational method to leverage the multi-modal readouts to predict TF binding events in cis-regulatory elements, which we then use to build GRNs that underlie cell fate transitions. Finally, we show that these models can be used to generate *in silico* predictions of the effect of TF perturbations.

## Results

### Simultaneous profiling of RNA expression and chromatin accessibility during mouse early organogenesis at single-cell resolution

We employed the 10× Multiome technology to profile RNA expression and chromatin accessibility from single nuclei collected between E7.5 and E8.75 (**Figure 1a**). A total of 61,781 cells passed quality control for both data modalities, with a median detection of 4,322 genes expressed per cell and a median of 29,710 ATAC fragments per cell (**Figure S1**). Cell types were assigned by mapping the RNA expression profiles to a reference atlas from similar stages (Pijuan-Sala et al., 2019) (**Figure 1b-c**, **Figure S2**). To evaluate the cell type assignments we performed multi-modal dimensionality reduction with MOFA+ (Argelaguet et al., 2020), revealing that both molecular layers contain sufficient information to distinguish cell type identities (**Figure 1c**). Similar results are obtained when applying dimensionality reduction to single data modalities. To further validate the measurements obtained from both data modalities, we compared the RNA expression and chromatin accessibility profiles with published data sets profiled with scRNA-seq (E7.5 to E8.5 embryos)(Pijuan-Sala et al., 2019) and snATAC-seq (E8.25 embryos)(Pijuan-Sala et al., 2020). Despite differences in the technology and in the molecular input (i.e. whole cell versus single nuclei in the case of RNA expression) we observe close agreements in both gene expression (**Figure S3**) and chromatin accessibility measurements (**Figure S4**).

**Figure 1:**
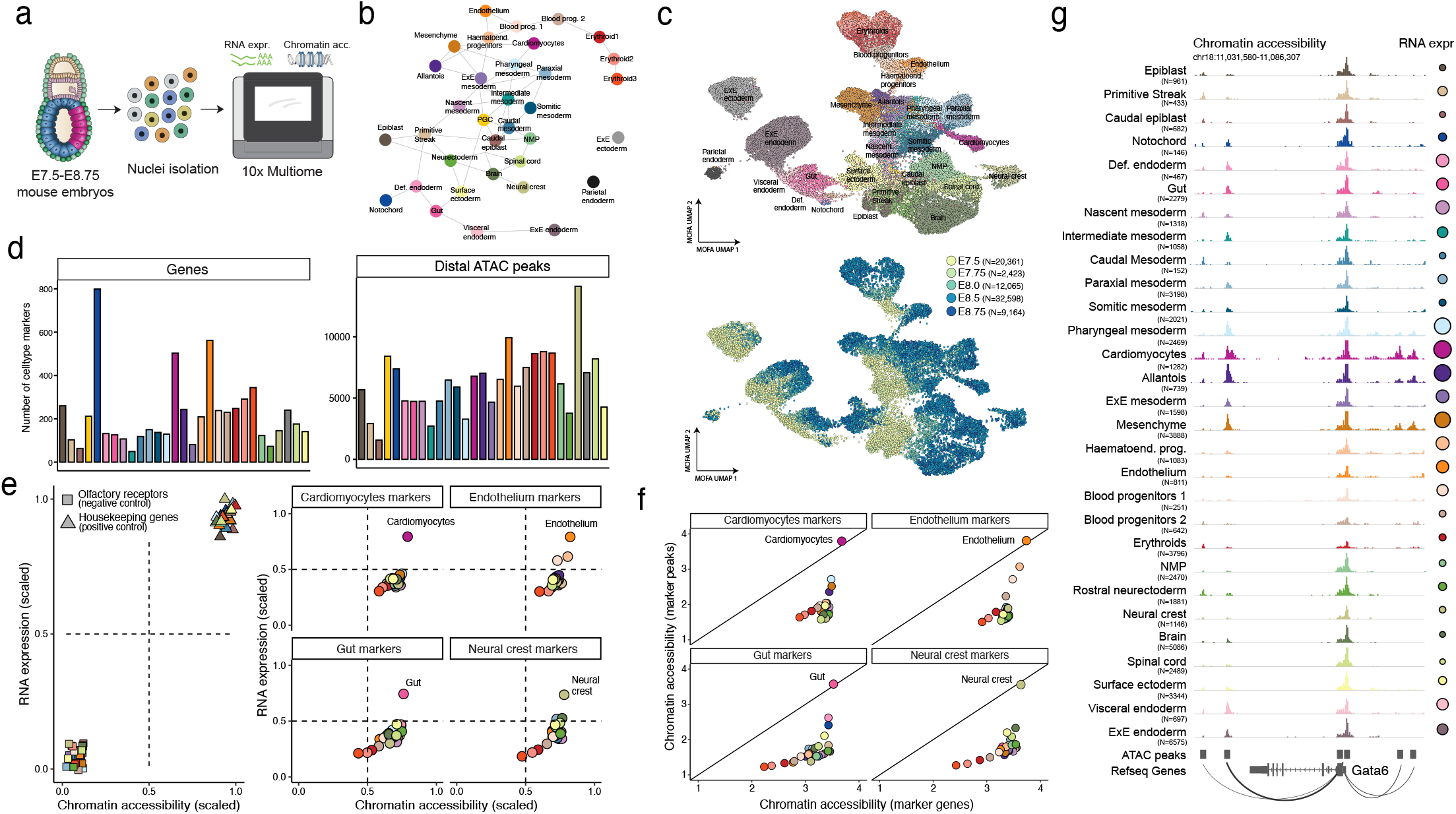
Simultaneous profiling of RNA expression and chromatin accessibility from single cells during mouse early organo-genesis. (a) Schematic display of the experimental design. Mouse embryos are dissociated into single cells then lysed to extract nuclei which are processed for simultaneous snATAC and snRNA-seq from the same cell using the 10× Multiome protocol. (b) Partition-based graph abstraction (PAGA) (Wolf et al., 2019) of the reference atlas (Pijuan-Sala et al., 2019), where each node corresponds to a different cell type. Cell types are coloured as per (Pijuan-Sala et al., 2019). (c) Multi-modal dimensionality reduction using MOFA, followed by UMAP (Argelaguet et al., 2020). Cells are coloured by cell type (top, see (b) for key) and stage (bottom). (d) Number of marker genes (left) and marker peaks (right) per cell type. See (b) for cell type colour key. (e) RNA expression and promoter chromatin accessibility values of different gene sets quantified separately for each cell type. The left panel shows olfactory receptors (negative control, non-expressed genes with closed chromatin) and housekeeping genes (positive control, highly expressed genes with open chromatin). The right panel shows different gene sets of cell type marker genes. Each dot corresponds to a pseudobulk cell type, coloured as in (b). Note that RNA expression and chromatin accessibility values are quantified as an average across all genes from each gene set. (f) Chromatin accessibility values of cell type marker genes (x-axis) and marker distal ATAC peaks from the same cell type (y-axis). Each panel shows gene and peak sets for different cell types. The diagonal line shows the values where both promoter and peak chromatin accessibility values are identical. Quantification of chromatin accessibility is done as in (e). (g) Genome browser snapshot of the *Gata6* locus. Each track displays pseudobulk ATAC-seq signal for a given cell type. Note the dynamic patterns of distal regulatory regions both upstream and downstream of the gene, compared to the uniformly open promoter region.

### A catalogue of cis-regulatory elements

To define open chromatin regions that represent putative cis-regulatory elements we performed peak calling on the snATAC-seq data using the ArchR pipeline (Granja et al., 2021). Briefly, peaks are defined by an iterative overlapping strategy where cells are aggregated by cell type into pseudo-bulk replicates. This approach has been shown to optimally preserve cell type-specific peaks (Granja et al., 2021). We obtained a total of 192,251 ATAC peaks, which we classified into four groups depending on their genomic location: Promoter (16.92%), Exonic (5.77%), Intronic (41.57%) and Intergenic (35.75%) (**Figure S5a-b**). 81% of peaks display differential accessibility in at least one cell type comparison (**Methods**). 69% of peaks were assigned to genes based on genomic proximity (less than 50kb from the gene body), with an average of ~20 peaks linked to a gene and an average of ~2.3 genes associated to a peak (**Figure S5c-d**). ~35% of peak-to-gene associations displayed significant positive correlation with the RNA expression levels of at least one of the proximal genes, whereas ~11% displayed a negative correlation (**Figure S5e-f**).

### Molecular characterisation of lineage-specific cis-regulatory elements

Next, we sought to characterise the transcriptomic and epigenetic variability of lineage-defining genes. We used the pairwise differential RNA expression results between cell types to define cell type-specific upregulated marker genes (**Figure 1d left, Methods**). Then, we quantified the average RNA expression and chromatin accessibility (at promoter regions) for each class of marker genes and each cell type (**Figure 1e right, Figure S6**). As a positive control, we performed the same quantification for a set of canonical housekeeping genes, which are constitutively expressed and have an open chromatin profile. As a negative control, we included a set of olfactory receptors genes, which are not expressed until later in development and display a closed chromatin profile (**Figure 1e left, Figure S6**). In marker genes, we observe the highest levels of expression and chromatin accessibility in the cell types that they mark, as expected. In all other cell types expression of these marker genes is still detected but at reduced levels. Promoter accessibility is also lower for marker genes in the cell types that they mark, however the differences are much less pronounced than for gene expression (**Figure 1e**). This suggests that promoter accessibility may have a limited function in driving differences in gene expression across cell types. Then, we asked whether cis-regulatory elements that are distal to promoter regions (Intronic and Intergenic peak sets) also display the same behaviour. We defined cell type-specific marker peaks by performing pairwise differential accessibility analysis (**Figure 1d right, Methods**), and then compared the average chromatin accessibility at promoter regions of marker genes versus marker peaks (**Figure 1f**). We find distal cis-regulatory elements to be more dynamic, with accessibility levels similar to promoters in the cell types where they become active, but much lower accessibility in the cell types where they are not active (**Figure 1f**). Consistent with previous reports (Argelaguet et al., 2019; Cusanovich et al., 2018), our results indicate a more prominent role of distal regulatory regions in cell fate decisions. A representative example is the *Gata6* locus shown in (**Figure 1g**). This gene is expressed in late mesodermal cell types, including Cardiomyocytes, Pharyngeal mesoderm and Allantois. However, the promoter region is homogeneously open across all cell types, whereas three regulatory regions located within 50 kilobases of the gene body gain accessibility exclusively in the cell types where *Gata6* is expressed. Other representative examples are shown in **Figure S6**.

### Multi-modal prediction of transcription factor binding sites

Cell fate decisions are molecularly characterised by changes in GRNs orchestrated by the interaction between TFs and their target genes (Levine and Davidson, 2005). Nevertheless, limited knowledge of TF binding exists for early embryonic development. First, experimental methods such as ChIP-seq or CUT&RUN require large numbers of cells to accurately profile TF binding events making it challenging to apply to embryos (Skene and Henikoff, 2017). Second, the success of the experiments depend on properties of available antibodies and on the properties of the TF itself, making it unrealistic to profile even a fraction of all transcription factors in the genome (Lambert et al., 2018; Park, 2009). Current methods for ATAC-seq data analysis link TFs to regulatory regions by the presence of TF motifs (Castro-Mondragon et al., 2021; Schep et al., 2017; Weirauch et al., 2014). This approach can be successful for some TFs that display non-redundant DNA motifs with high sequence specificity, but it has important shortcomings. First, the presence of a TF motif does not guarantee the existence of an active binding site (Wang et al., 2012). Second, a large fraction of TFs belong to families that share the same motif, even when having different functions and expression patterns. Representative examples are the GATA, HOX and the FOX family of transcription factors (**Figure S7a-b**). As a result of these issues, it is extremely challenging to link TFs to regulatory elements when exclusively using a combination of genomic and epigenomic information. This can be illustrated by the large number of TF motifs that can be identified within each ATAC peak (**Figure S7c-d**). To address this, we developed a computational approach that integrates genomic, epigenomic and transcriptomic information to predict TF binding events.

Intuitively, we consider an ATAC peak *i* to be a putative binding site for TF *j* if it contains the *j* motif and its chromatin accessibility is correlated with the RNA expression of the TF (**Figure 2a**). We combine three metrics (motif score, average chromatin accessibility and correlation) to devise a quantitative *in silico* binding score for each combination of TF and ATAC peak (**Methods**). Note that our approach is unsupervised and does not require ChIP-seq data as input. This stands in contrast with other approaches that have been proposed to predict TF binding from multi-omics data, which employ supervised models that require labelled training data from ChIP-seq experiments (Avsec et al., 2021; Karimzadeh and Hoffman, 2019). We will refer to this approach as *in silico* ChIP-seq. The number of predicted binding sites for each TF is a function of the minimum score threshold, which ranges from 0 to 1 after scaling (**Figure 2b**). Notably, the incorporation of RNA expression massively reduces the amount of predicted binding sites for each TF as well as the amount of TFs that can be linked to each regulatory element (**Figure 2c-d**).

**Figure 2:**
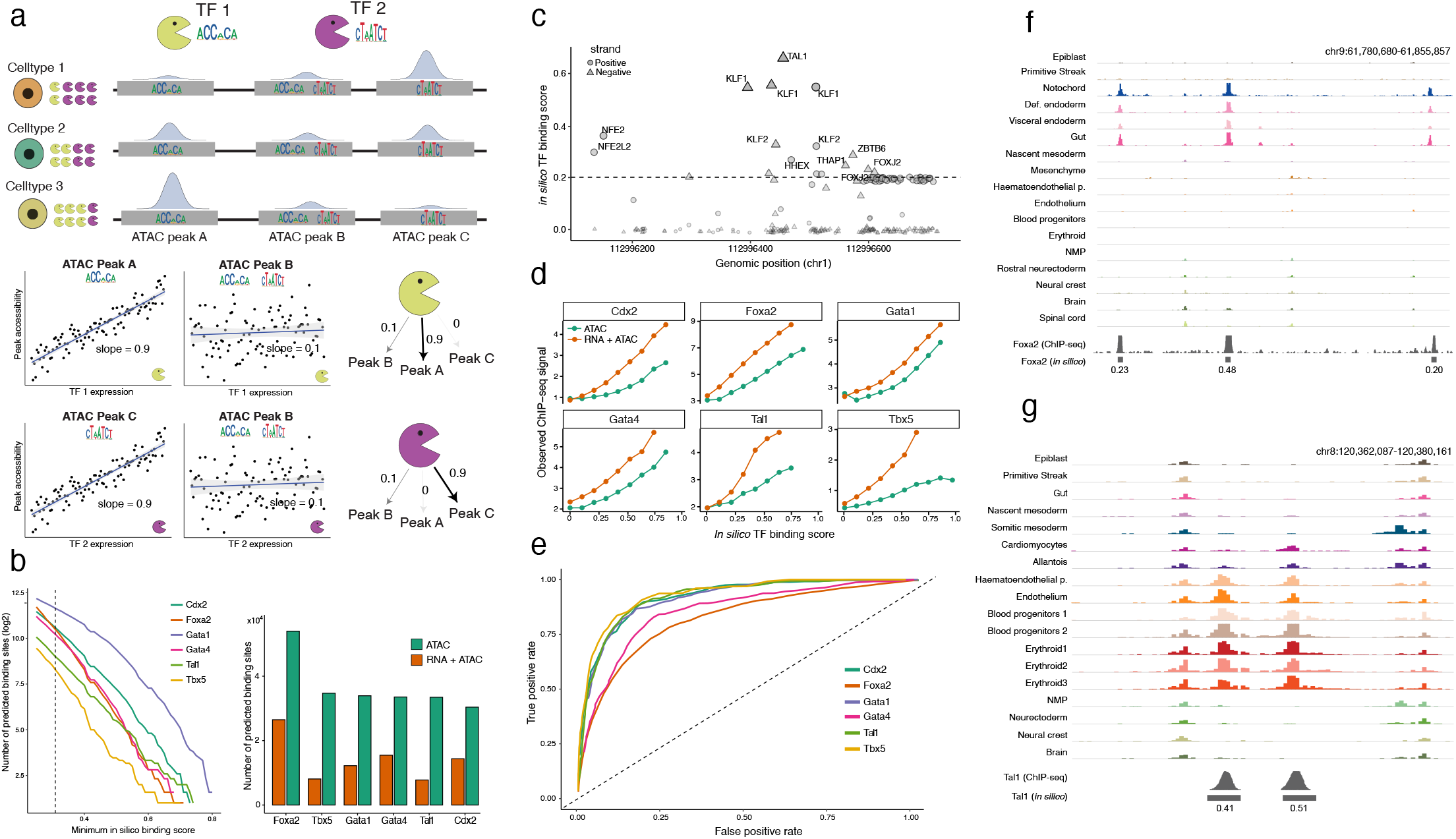
*In silico* ChIP-seq: leveraging multi-modal information to predict transcription factor binding sites. (a) Schematic of the *in silico* ChIP-seq methodology. Consider two different TFs (Pacmans), each one with different DNA binding preferences encoded in the form of different position-specific weight matrices; and three cis-regulatory elements represented as ATAC peaks (grey boxes), each one containing different instances of the TF motifs. Each row displays a different cell (or metacell or cell type, depending on the level of data aggregation). Each cell type is associated with different values of TF RNA expression (see changes in Pacman abundance) and chromatin accessibility of the cis-regulatory elements (see changes in the density histogram). The *in silico* ChIP-seq model exploits the correlation between TF RNA expression and the chromatin accessibility of the ATAC peaks that contain at least one instance of its TF motif to derive a quantitative TF binding score. In the schematic Peak A contains the TF 1 motif, and its accessibility correlates with the RNA expression of TF 1, thus leading to a high TF binding score. Peak B also contains the TF 1 motif, but its accessibility correlates poorly with the TF’s RNA expression, which leads to a non-zero but low TF binding score. Peak C does not contain the TF 1 motif, which leads to a zero TF binding score. (b) Left: the number of predicted binding sites for 6 representative TFs as a function of the minimum *in silico* TF binding score. Dashed line indicates the minimum score used in subsequent analyses. Right: Bar plots showing the number of predicted binding sites in the *in silico* ChIP-seq model when incorporating the RNA expression (orange) versus just using ATAC information (green). (c) A representative instance of an ATAC peak highlighting the large number of TF motifs contained within a 600bp locus. Shown are the positions of all TF motifs within the ATAC peak (x-axis) against the *in silico* ChIP-seq score (y-axis). Note that only a subset of TF motif instances display high *in silico* ChIP-seq score. The dashed line indicates the cutoff used to determine a putative binding site, as in (b). (d) Comparison of *in silico* TF binding scores (x-axis) versus experimental ChIP-seq signal (y-axis), using the same 6 TFs as in (b). Orange line displays scores derived from the *in silico* ChIP-seq model, whereas the green line displays scores derived when just using ATAC-seq information (i.e. omitting the TF RNA expression from the model). Scores were binned from 0 to 1 in intervals of 0.1,and each dot corresponds to the average value across all cis-regulatory regions from the interval. ChIP-seq datasets are all derived from publicly available data sets that most closely resemble mouse embryos at the gastrulation and organogenesis stage (Supplementary Table 1). (e) Receiver Operating Characteristic (ROC) curves comparing the predicted TF binding sites vs the real TF binding sites (inferred from peak calling on the experimental ChIP-seq data). (f) Genome browser snapshot displaying *Foxa2* binding sites. Each track displays pseudobulk ATAC-seq signal for a given celltype. The experimental ChIP-seq values are shown in the bottom, together with the *in silico* TF binding scores for the ATAC peaks that have a TF binding score higher than 0.20 (same threshold as in (b)). (g) Same as in (f) but displaying *Tal1* binding sites.

To validate the *in silico* ChIP-seq library, we used publicly available ChIP-seq experiments for a set of TFs that are known to play key roles during mouse gastrulation and early organogenesis, and defined this as the ground truth for TF binding events. Due to the limited availability of *in vivo* ChIP-seq datasets, we had to rely on *in vitro* models that more closely resemble the gastrulating embryo (**Supplementary Table 1**). Yet, we observe remarkable agreement between the *in silico* TF binding scores and the observed ChIP-seq signal (**Figure 2d-e**). Worse agreement is obtained when excluding the transcriptomic information from the model (**Figure 2d**). Representative examples of TF binding predictions are shown alongside ChIP-seq data in **Figure 2f-g**. Interestingly, for all TFs we benchmarked, the consistency with ChIP-seq measurements exclusively holds true for ATAC peaks that are positively correlated with TF expression (**Figure S8**), which is consistent with these TFs acting as chromatin activators. Our approach also predicts repressive interactions with chromatin (not to be confused with transcriptional repression of target genes, as we will discuss below). Chromatin repressors are known to be important for gene regulation, and they generally involve the recruitment of chromatin remodelers, including histone modifiers, to turn chromatin from an open to a closed state (Berest et al., 2019; Gaston and Jayaraman, 2003; Iurlaro et al., 2021; Janssens et al., 2022; Lambert et al., 2018). However, insufficient ChIP-seq data exists for chromatin repressors in the context of embryonic development, thus limiting our benchmark. In consequence, we only consider activatory links between TFs and regulatory regions for downstream analyses. We refer the reader to the Supplementary Information for a more detailed discussion on the methodology, benchmark, limitations of the method and comparison to related approaches.

### Quantification of cell type-specific transcription factor chromatin activities using chromVAR-Multiome

The output of the *in silico* ChIP-seq model is a matrix of TF binding scores for each cis-regulatory element. It does not directly provide a quantification of the TF activities for each sample (where samples can correspond to cells, metacells or cell type, depending on the chosen data resolution, see discussion in the **Supplementary Information**). The most popular method to quantify TF activities per sample using chromatin accessibility data is chromVAR (Schep et al., 2017). Briefly, this method computes, for each sample and TF motif, a z-score that measures the difference between the total number of fragments that map to motif-containing peaks and the expected number of fragments (**Figure 3a**). While useful when only having access to chromatin accessibility data, chromVAR scores are often not representative of true TF activities, mainly because accessible DNA motifs are not always good proxies for actual TF binding events. This can be illustrated with the Fox family of transcription factors, which all share a similar DNA motif, but nevertheless have different roles during mouse gastrulation: whereas *Foxb1* is a pioneer TF in the ectodermal lineage (Labosky et al., 1997)*, Foxc2* is active in the mesodermal lineage (Wilm et al., 2004). Their distinct roles are evidenced by the different cell types and spatial locations where they are expressed in the embryo (Lohoff et al., 2022) (**Methods, Figure 3b-c**). Yet, due to their motif similarity, the chromVAR scores of these two TFs are indistinguishable (**Figure 3b-c**). Here, we modified the chromVAR algorithm to use the putative TF binding sites from the *in silico* ChIP-seq library, instead of all TF motif instances. We refer to this approach as chromVAR-Multiome, and the resulting values as TF activity scores.

**Figure 3:**
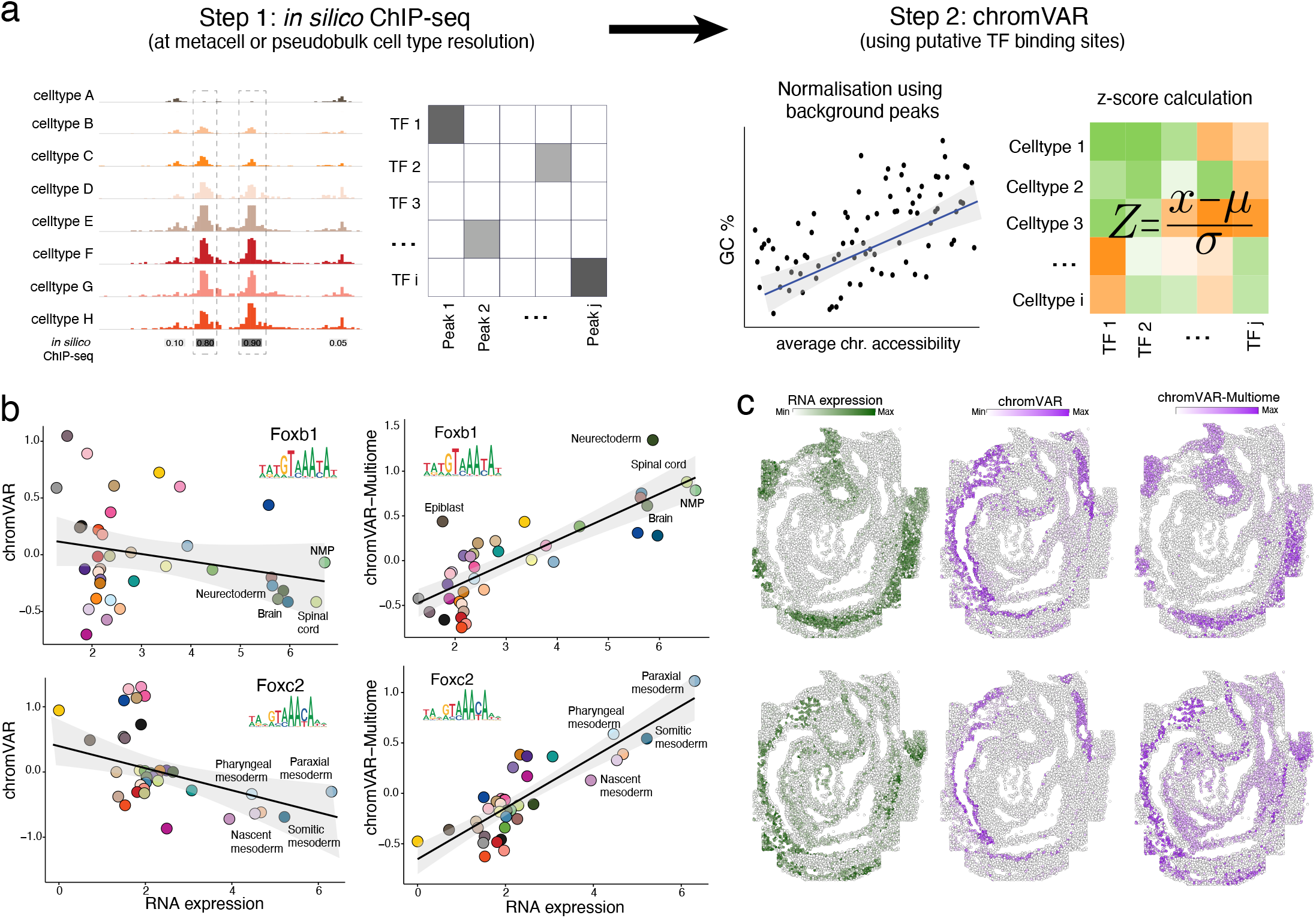
Quantification of celltype-specific transcription factor chromatin activities with chromVAR-Multiome. (a) Schematic of chromVAR-Multiome. In the first step, the *in silico* ChIP-seq method is used to calculate putative transcription factor (TF) binding sites. This results in a matrix of TF binding scores for each combination of TF and ATAC peak. In the second step, we calculate celltype-specific TF activities using the chromVAR algorithm (Schep et al, 2017) but replacing the default input of all motif-containing ATAC peaks with putative TF binding events. The chromVAR-Multiome algorithm yields z-scores for each combination of TF and cell type (or metacell, depending on the chosen data resolution). (b) Comparison of chromVAR and chromVAR-Multiome for quantification of TF chromatin activities for Foxb1 (top) and Foxc1 (bottom). Scatter plots display the TF’s RNA expression (x-axis) and the chromatin accessibility of target regions (y-axis), quantified at the pseudobulk level using chromVAR (left) or chromVAR-Multiome (right). Each dot corresponds to a different cell type. (c) Spatially-resolved RNA expression (imputed values from Lohoff et al, 2021, coloured in green) and TF chromatin activity (coloured in purple) for Foxb1 (top) and Foxc1 (bottom), quantified using chromVAR or chromVAR-Multiome. Note that spatially-resolved TF chromatin activity values are inferred by mapping the 10× Multiome cells onto the spatial transcriptomic data set.

Overall we find that chromVAR-Multiome yields TF activity scores that are more consistent with the known expression patterns of the corresponding TFs (**Figure 3b**), albeit with some exceptions. By exploring the residuals of a linear model linking chromVAR-Multiome scores with TF RNA expression one can identify outlier cell types that are candidates for epigenetic priming (**Figure S9**). An example is *Foxb1*: while high levels of RNA expression and TF activity are observed in mature ectodermal lineages, including Spinal cord and Brain, Epiblast cells display high *Foxb1* activity despite the TF not being expressed (**Figure S9**). Notably, this observation is consistent with our previous work (Argelaguet et al., 2019), where we proposed that pluripotent epiblast cells are epigenetically primed for neuroectoderm differentiation via epigenetic priming of cis-regulatory elements.

### A catalogue of cell type-specific transcription factor activities in mouse early organogenesis

Next, we used the chromVAR-Multiome scores to perform pairwise differential analysis between cell types and parse the results to quantify TF markers for each cell type (**Methods**) (**Figure 4a-b**). Reassuringly, using this approach we recover canonical TF markers for a variety of cell types, including Foxa2 and Sox17 for endodermal cell types; Mesp11/2 and Mixl1 for the Primitive Streak and mesodermal cell types; Sox2 and Rfx4 for ectodermal cell types; Tbx5 and Nkx2-5 for Cardiomyocytes; Runx1 and Tal1 for Blood progenitors and Erythroids (**Figure 4c**). Notably, the resolution of the data enables us to provide quantifications of TF activities for cell types that are challenging to study due the low cell numbers and difficult cell isolation, including Primordial Germ Cells (PGCs) and Neural crest cells (**Figure 4d**). For the Neural crest, we recover many TFs that have been previously associated with Neural crest identity in different species: Pax7, Foxd3, Tfap2a, Tfap2b, Sox10, Sox5, Ets1, Nr2f1 and Mef2c (**Figure S10**, **Supplementary Table 2**). For example, Tfap2a has been shown to be essential for Neural crest specification in Xenopus embryos (de Crozé et al., 2011). In mice, disruption of the *Tfap2a* gene results in craniofacial malformations and embryonic lethality(Schorle et al., 1996). In humans, missense mutations in the corresponding orthologous gene results in branchio-oculo-facial syndrome, which is also characterised by craniofacial abnormalities (Milunsky et al., 2008). For PGCs we also recover TFs described to be important for PGC specification in mice, including Prdm1 (also called Blimp-1), Esrrb and Pou5f1 (also called Oct4) (**Figure S10**, **Supplementary Table 2**). For example, Blimp-1 has been shown to be essential for the repression of the somatic programme upon PGC specification(Ohinata et al., 2005). In addition, we also predict several TFs with unknown roles in PGC formation that could be suitable candidates for further characterisation, including Ybx2, Bbx and Klf8 (**Figure S10**).

**Figure 4:**
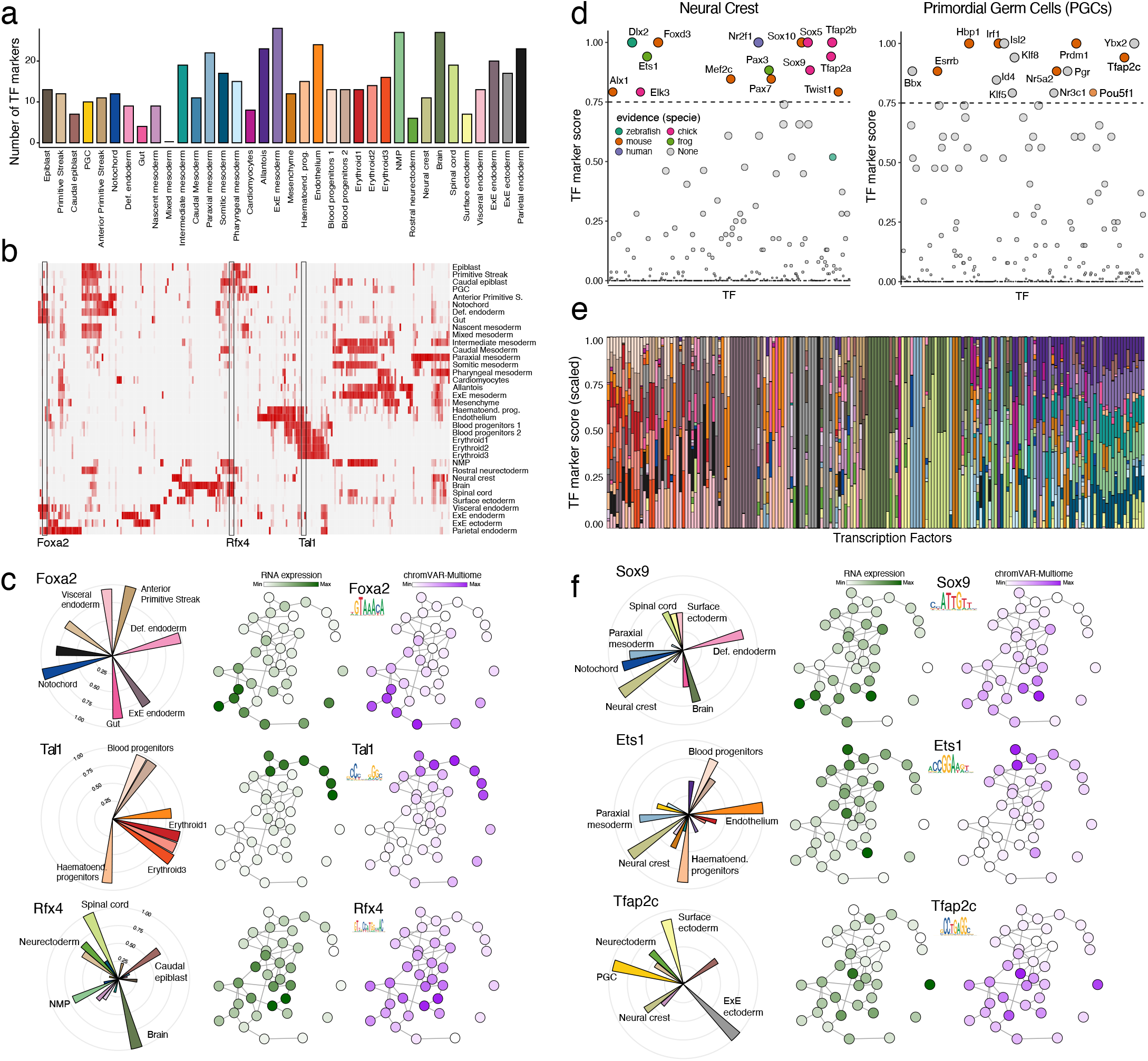
A catalogue of cell type-specific transcription factor activities during mouse early organogenesis reveals widespread pleiotropy. (a) Barplot displaying the number of TF markers per cell type. TF markers are inferred using the TF activity scores, which results from performing differential analysis with the chromVAR-Multiome values (Methods). The higher the score for TF i in celltype j, the more active this TF is predicted to be in cell type j, with a minimum score of 0 and a maximum score of 1. (b) Heatmap displaying TF activity scores for each celltype (rows) and each TF (column). (c) Left: polar plots displaying the celltype TF activity scores for three different TFs: Foxa2 (top), Tal1 (middle) and Rfx4 (bottom). Right: PAGA representation of the transcriptomic atlas as in Figure 1b for the three same TFs, with each node coloured by the RNA expression of the TF (green) and the corresponding chromVAR-Multiome score (purple). (d) Dot plots displaying the TF activity scores for all TFs in Neural Crest cells (left) and PGCs (right). TFs with the highest TF activity score are labelled and coloured to indicate whether a known function has been reported and in which species the evidence was obtained (Supplementary Table 1). (e) Stacked bar plots displaying the (scaled) TF activity scores for each combination of TF and cell types. Each column corresponds to a TF. (f) as (c) but for TFs that display a pleiotropic effect (i.e. they are active across distinct cell types).

Interestingly, visualisation of TF activities across all cell types reveals that (1) cell types are defined by a combinatorial activity of multiple TFs and (2) most TFs are active across multiple cell types (**Figure 4e**). The first observation can be illustrated with the Neural crest: Of the canonical TFs shown in **Figure 4f**, none are uniquely active in the Neural crest, with the exception of Dlx2 and Sox10. The second observation sometimes arises from the hierarchical nature of lineage specification (such as Pax7 being active in multiple ectodermal-derived cell types, Foxa2 in all endodermal-derived cell types and Tal1 in all cell types that are linked to blood formation). However, in other cases we observe the same TF active in cell types from different germ layer origins, thus suggesting widespread pleiotropic activity where TFs define cellular identities via combinatorial context-dependent activity (Reiter et al., 2017; Spitz and Furlong, 2012). Representative examples are Sox9, active in the Neural crest, Brain, Definitive endoderm, and Notochord; Tfap2c, active in the Neural crest, ExE ectoderm and PGCs; and Ets1, active in Neural crest, Endothelium and Blood Progenitors (**Figure 4f**).

### Mapping the transcription factor regulatory network that underlies differentiation of neuromesodermal progenitors

In the previous section, we used the chromVAR-Multiome scores to generate a catalogue of TF activities linked to cell types. In this approach, however, we ignored interactions between TFs. Next, we sought to quantify interactions between TFs by inferring gene regulatory networks (GRNs) and connecting them to continuous cellular trajectories.

We employed a multi-step algorithm to infer GRNs (**Methods, Figure S11**). First, we subset cells of interest and infer metacells (Persad et al., 2022), with the goal of achieving a resolution that retains the cellular heterogeneity while overcoming the sparsity issues of single-cell data. Second, we used the *in silico* ChIP-seq method to link TFs to cis-regulatory elements. Third, we linked cis-regulatory elements to potential target genes by genomic proximity (here a conservative maximum distance of 50kb), which is a reasonable approximation in the absence of 3D chromatin contact information (Janssens et al., 2022; Kamal et al., 2021). This results in a directed network where each parent node corresponds to a TF, and each child node corresponds to a target gene. Finally, following the approach of (Kamimoto et al., 2020), we estimated the weights of the edges by fitting a linear regression model of target gene expression as a function of the parent TF’s expression. Importantly, while our benchmark of the *in silico* ChIP-seq does not support the inclusion of repressive links between TFs and cis-regulatory elements, evidence exists that TFs can repress the expression of target genes (Gaston and Jayaraman, 2003; Liang et al., 2017). Thus, in the GRN model we allowed for negative associations between TF expression and target gene expression.

Here we applied the GRN methodology described above to study the TF regulatory network underlying differentiation of Neuromesodermal progenitors (NMPs). Briefly, NMPs are a population of bipotent stem cells that fuel axial elongation by simultaneously giving rise to Spinal cord cells, an ectodermal cell type, as well as posterior somites, a mesodermal cell type (Sambasivan and Steventon, 2020) (**Figure 5a**). Molecularly, NMPs are characterised by the co-expression of the mesodermal factor Brachyury and the neural factor Sox2 (Henrique et al., 2015), but studies have suggested that these are just two players of a complex regulatory landscape (Gouti et al., 2017). Consistently, besides Sox2 and Brachyury, we identify a network of 33 additional TFs with 379 activatory and 48 repressive interactions, respectively (**Figure 5b**), Notably, we find the homeobox Cdx and Hox TFs display the highest centrality of the network by establishing an activatory self-regulatory loop that sustains NMP identity (**Figure 5c,d**). This observation agrees with studies that showed that all three Cdx genes contribute additively to axial elongation and the development of posterior embryonic structures, with the most important one being Cdx2 <u>(Chawengsaksophak et al. 2004; van Rooijen et al. 2012; Metzis et al. 2018)</u>. To further validate the predicted interaction between Cdx and Hox genes, we used ChIP-seq data for Cdx2 profiled in Epiblast Stem Cells exposed to Wnt and Fgf signalling, which induces posterior axis elongation and generates cells that resemble NMPs (Amin et al., 2016). Consistent with the inferred GRN, we find widespread binding of Cdx2 within the *Hoxb* cluster of genes (**Figure 5e**). This interaction between Cdx and Hox genes also agrees with *in vitro* studies that described the upregulation of posterior Hox genes in NMP-like cells upon induction of Cdx factors (Amin et al., 2016; Neijts et al., 2016). Interestingly, in addition to its role as transcriptional activator of Hox genes, we also find that Cdx2 displays a pleiotropic role by repressing TFs that direct the transition to Somitic mesoderm (Foxc2, Brachyury, Meox1) and Spinal cord (Pax6) (**Figure 5f and Figure S12**).

**Figure 5:**
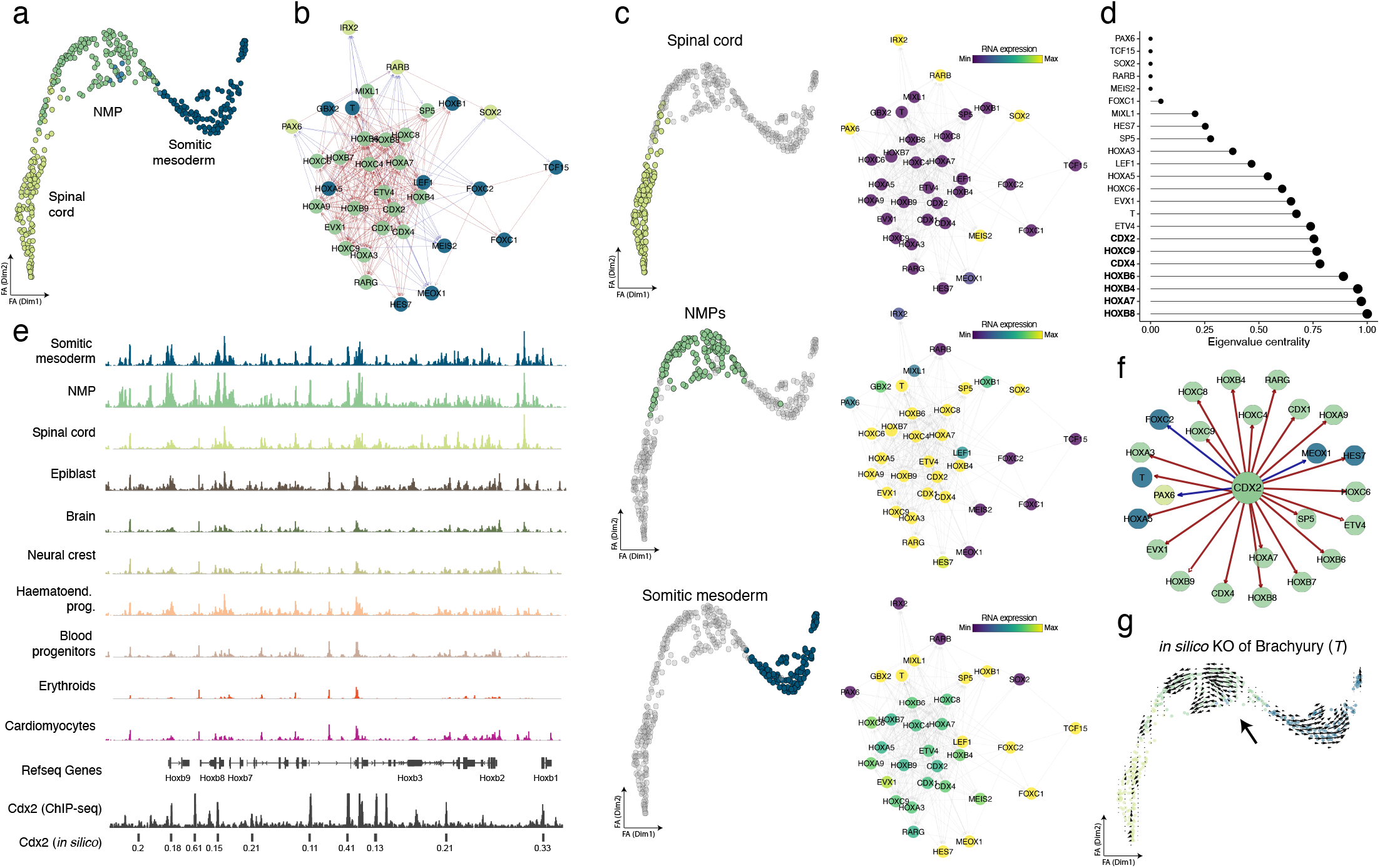
Characterisation of the transcription factor regulatory network underlying Neuromesodermal progenitors. (a) Force-atlas layout of the NMP differentiation trajectory. Each dot corresponds to a metacell, coloured by cell type identity. (b) TF regulatory network inferred using the NMP trajectory. Each node corresponds to a TF, coloured by the cell type where the TF displays the highest expression. Edges denote regulatory relationships: red edges represent activatory relationships (the expression of the parent node is positively correlated with the expression of the child node), whereas blue edges represent repressive relationships (the expression of the parent node is negatively correlated with the expression of the child node). (c) Left: same layout as in (a) but highlighting each of the three cell types of the trajectory: Spinal cord (top), NMP (middle) or Somitic mesoderm (bottom). Right: Same TF regulatory network as in (b), but nodes are coloured based on the average expression of the TF in each of the three cell types of the trajectory: Spinal cord (top), NMP (middle) or Somitic mesoderm (bottom). For clarity, we increased the transparency of edges. (d) Eigenvalue centrality for each TF in the network. (e) Genome browser snapshot of the *Hoxb* loci. Each track displays pseudobulk ATAC-seq signal for a given celltype. Shown in the bottom is the *in silico* ChIP-seq predictions for Cdx2 and the experimental ChIP-seq signal for Cdx2 profiled in NMP-like cells. (f) Regulatory connections between Cdx2 and downstream TFs. As in (b), nodes are coloured by the cell type where the TF displays the highest expression. (g) *in silico* knock-out of Brachyury using CellOracle (Kamimoto et al, 2021). Shown is the same layout as in (a), with arrows displaying the predicted changes in cell state for different parts of the trajectory when knocking out Brachyury.

### *Brachyury* controls the differentiation of Neuromesodermal progenitors to Somitic mesoderm by epigenetic priming cis-regulatory elements

Our results above indicate that Cdx and Hox factors sustain the bipotent NMP identity but can also act as a fate switch by repressing TFs that direct the transition to Somitic mesoderm, most notably Brachyury. To validate whether this role of Brachyury is captured by the gene regulatory networks, we employed the GRN of NMP differentiation to predict the consequences of Brachyury knock-out using the CellOracle framework (Kamimoto et al., 2020) (**Methods**). Briefly, CellOracle enables network configurations to be interrogated *in silico* by simulating the effects of TF perturbations. By leveraging the visualisation framework of RNA velocity, CellOracle can be used to predict how cell state shifts after perturbation of individual genes. Notably, we find that the *in silico* knock-out of Brachyury disrupts the transition from NMP to Somitic mesoderm (**Figure 5g**). Although this result was expected based on experimental evidence (Guibentif et al., 2021), it demonstrates how GRNs inferred from unperturbed single-cell multi-omics data have the potential to provide functional insights into cell fate transitions.

To further validate our predictions and obtain additional mechanistic insights, we generated Brachyury KO embryos by direct delivery of CRISPR/Cas9 as a ribonucleoprotein (RNP) complex via electroporation, targeting exon 3 of the *Brachyury* (*T*) gene in zygotes at one-cell stage (**Methods, Figure 6a**). Control embryos received Cas9 protein only. Embryos were transferred into pseudopregnant females and collected at E8.5 for 10× Multiome sequencing. In total, we obtained 6,797 cells from 3 embryos at E8.5 with a wildtype (WT) background and 6,572 cells from 7 embryos with a *Brachyury* KO background. Cell types were again annotated by mapping the RNA expression to the transcriptomic gastrulation atlas (**Figure 6b**).

**Figure 6:**
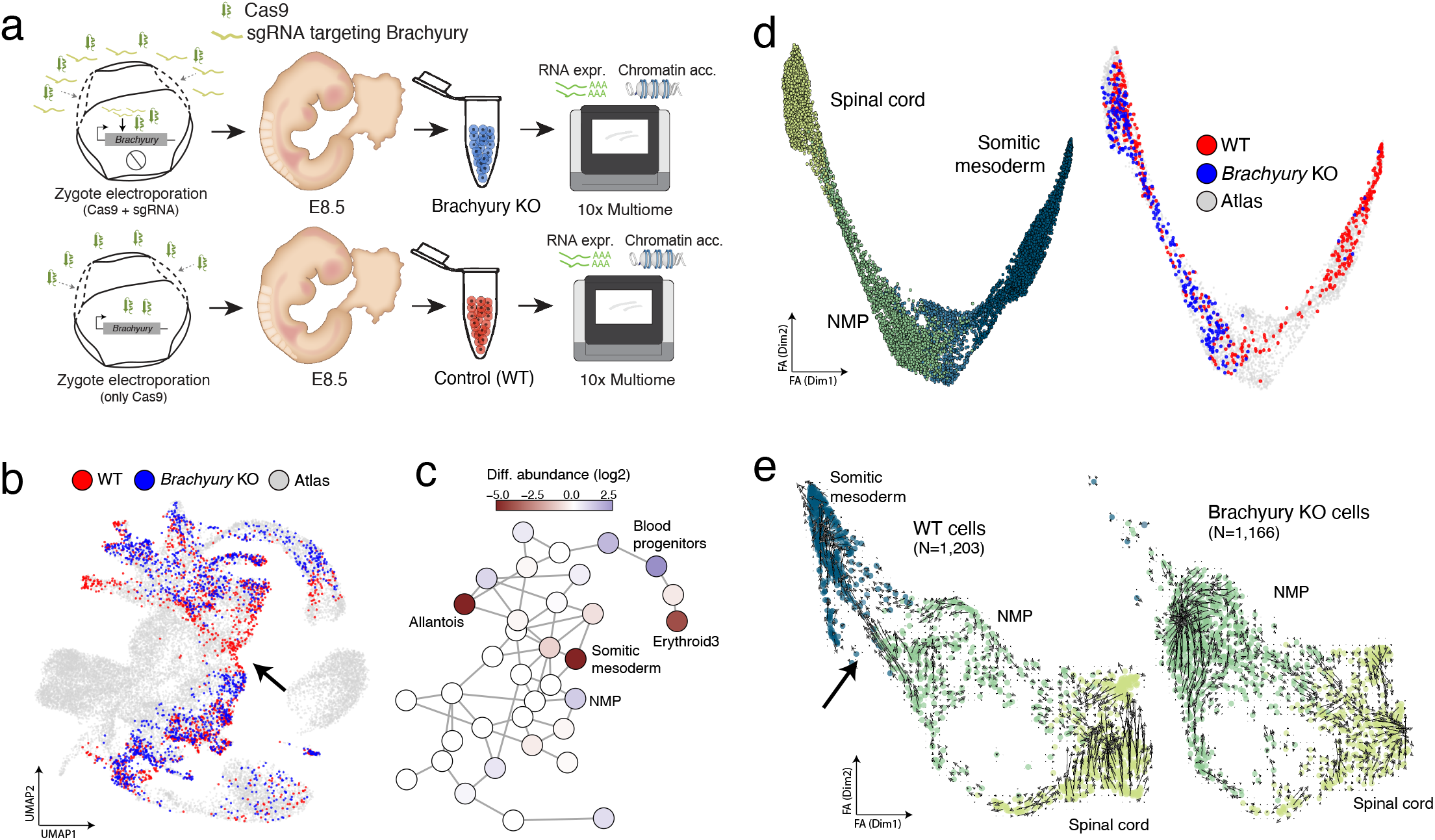
Brachyury is essential for the differentiation of neuromesodermal progenitors to somitic mesoderm. (a) Schematic showing the experimental design. We generated Brachyury KO embryos by electroporation of Cas9 protein and a single guide RNA (sgRNA) targeting the Brachyury gene (T). Control embryos received Cas9 but no sgRNA. Embryos were transferred into pseudopregnant females and collected at E8.5 for 10x Multiome sequencing. (b) Mapping WT and Brachyury KO cells to the reference atlas (Pijuan-Sala et al., 2019). Highlighted are cells in the reference dataset that are nearest neighbours to WT cells (red) or Brachyury KO cells (blue) in this experiment. (c) PAGA representation of the reference atlas, where each node corresponds to a cell type. Nodes are coloured by differences in cell type abundance between WT and Brachyury KO cells. Positive values indicate more abundance in the Brachyury KO, negative values indicate less abundance in the Brachyury KO. (d) Force-directed layout of the trajectory that connects Neuromesodermal Progenitor (NMP) cells to either Spinal cord or Somitic mesoderm, inferred using the reference atlas. Left: each cell is coloured by cell type. Right: mapping cells to the reference NMP trajectory. Highlighted are cells in the reference trajectory that are nearest neighbours to WT cells (red) or Brachyury KO cells (blue) in this experiment. (e) RNA velocity analysis of the NMP trajectory using scVelo (Bergen et al. 2020) on the 10x Multiome cells. Shown are WT cells (left) and Brachyury KO cells (right). The arrow highlights the trajectory from NMP to Somitic mesoderm that is present in WT cells but absent in Brachyury KO cells.

Consistent with our predictions and the results of (Gouti et al., 2017; Guibentif et al., 2021), we observe a relative underrepresentation of (posterior) Somitic mesoderm and Allantois cells in the *Brachyury* KO embryos, together with a relative overrepresentation of NMP cells (**Figure 6c**). No significant differences are observed in the abundance of Spinal cord cells, suggesting that the neural differentiation capacity of NMPs is not affected in the absence of Brachyury. Interestingly, we also observe defects in the Erythropoiesis trajectory (**Figure 6c**), suggesting pleiotropic effects of Brachyury across multiple developmental trajectories (Bruce and Winklbauer, 2020). To further explore the effect of the *Brachyury* KO in NMPs, we mapped the cells onto the NMP differentiation trajectory reconstructed from the transcriptomic reference atlas (Pijuan-Sala et al., 2019) (**Figure 6d**). Again, we find that WT cells map across the entire trajectory, but *Brachyury* KO cells map only onto the transition between NMP and Spinal cord. Additionally, RNA velocity analysis of these cells shows that WT NMP cells transition towards both Spinal cord and Somitic mesoderm fates, whereas in the *Brachyury* KO only the Spinal cord displays a coherent differentiation trajectory (**Figure 6e**).

Next, we performed differential accessibility analysis between WT and Brachyury KO NMP cells (**Methods**), and found 399 differentially accessible (DA) ATAC peaks. Notably, most of them (N=344) are open in WT cells but unable to open in Brachyury KO cells (**Figure 7a**). This set of DA peaks display enrichment for the T-box motif and a higher *in silico* TF binding score for Brachyury than non DA peaks (**Figure 7b-c**), hence indicating a direct regulation by Brachyury. Interestingly, some of these putative cis-regulatory elements are linked to mesodermal genes that become expressed in the Somitic mesoderm, including Tbx6, Mesp1 and Fgf4 (**Figure 7d-e**, **Figure S13**). Visualisation of the chromatin accessibility dynamics at these loci reveals that they attain the highest level of accessibility in the Somitic mesoderm cells, consistent with the increased expression of their target genes. Interestingly, however, we find these elements to be open in the progenitor NMP cells, before any expression of mesodermal genes is observed (**Figure 7d-e, Figure S13**). This behaviour is suggestive of epigenetic priming, whereby the chromatin of cis-regulatory elements becomes accessible before transcription of the target gene (Argelaguet et al., 2019; Ma et al., 2020). A representative example is shown in **Figure 7f**, which shows a cis-regulatory element that is targeted by Brachyury and is located upstream of *Mesp1*. This element becomes partially open in WT NMP cells, but not in *Brachyury* KO NMP cells, and attains its highest accessibility levels in WT Somitic mesoderm cells, while becoming closed in Spinal cord cells. Similar patterns can be observed for the cis-regulatory elements linked to *Tbx6* and *Fgf4* (**Figure S14)**. In conclusion, our results suggest that formation of posterior Somitic mesoderm is associated with Brachyury-driven epigenetic priming of cis-regulatory elements in NMP cells.

**Figure 7:**
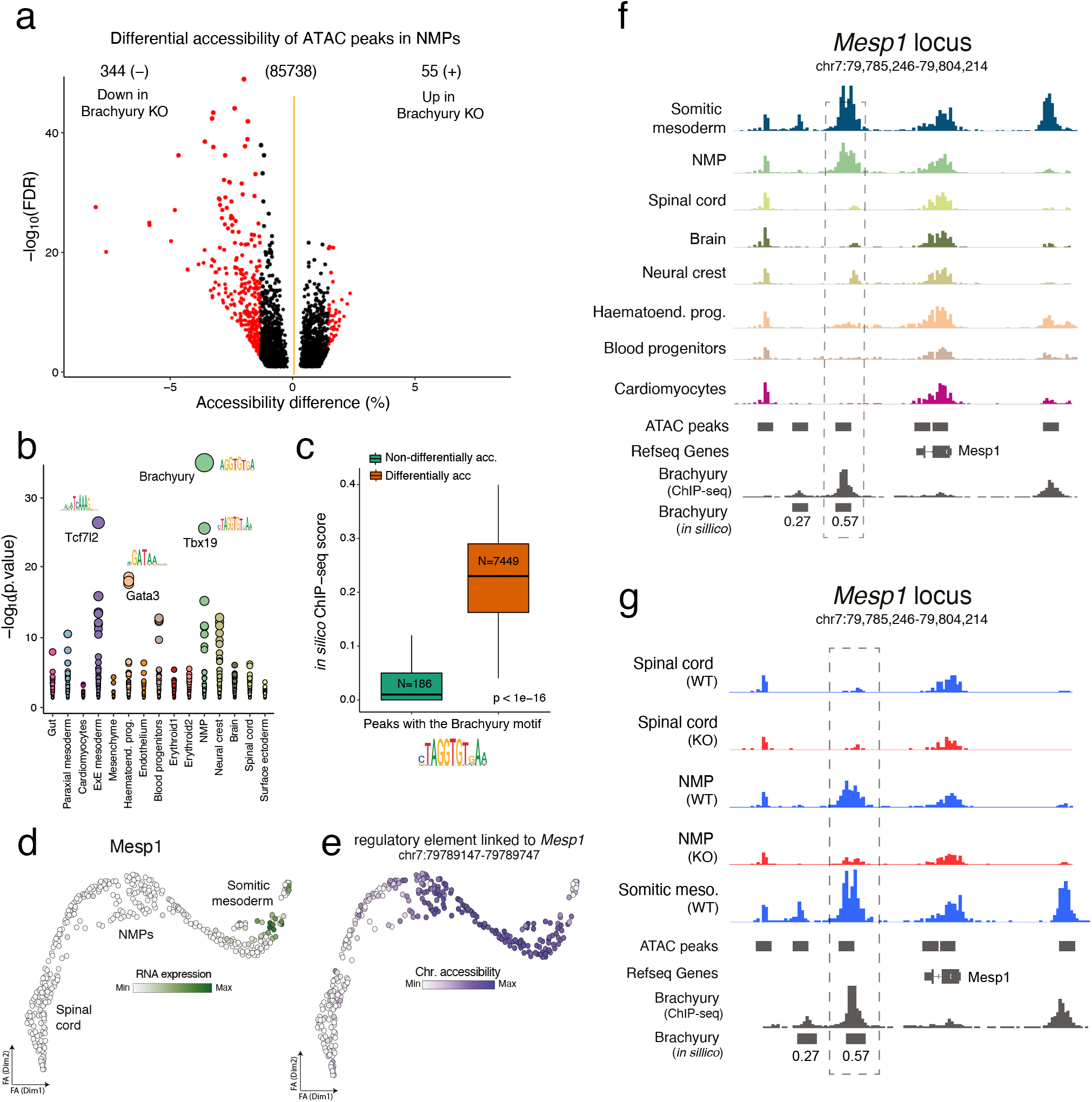
Brachyury controls the transition from neuromesodermal progenitors to somitic mesoderm by priming cis-regulatory elements. (a) Volcano plot displays differential accessible ATAC peaks between WT and Brachyury KO NMP cells. Coloured in red are ATAC peaks that pass statistical significance threshold. (b) TF motif enrichment analysis in differentially accessible peaks per cell type (x-axis). The y-axis displays the FDR-adjusted p-values of a Fisher exact test. Each dot corresponds to a different TF motif, coloured by the cell type where the differential accessibility analysis is performed. (c) Box plots display *in silico* TF binding scores for Brachyury within ATAC peaks that contain the Brachyury motif. ATAC peaks are split based on their differentially accessibility significance when comparing WT and Brachyury KO NMP cells. Note that the *in silico* ChIP-seq is inferred using metacells from the NMP trajectory from the reference atlas and does not include Brachyury KO cells. (d) Force-atlas layout of the NMP differentiation trajectory. Each dot corresponds to a metacell, coloured by the RNA expression of Mesp1. (e) Same layout as in (d), but metacells are coloured by the chromatin accessibility of the cis-regulatory element linked to *Mesp1*. (f) Genome browser snapshot of the chromatin accessibility signal around the *Mesp1* loci. Each track displays pseudobulk ATAC-seq signal for a given celltype from the reference atlas. Shown in the bottom is the experimental ChIP-seq signal for Brachyury profiled in Embryoid Bodies (Tosic et al 2019) and the corresponding *in silico* ChIP-seq predictions. Highlighted is a Brachyury-targeted regulatory element located downstream of *Mesp1*. (g) Genome browser snapshot of the same loci displayed in (f), but showing the ATAC-seq signal for a given celltype and genotype (WT in blue and Brachyury KO in red).

## Conclusion

We have generated a single-cell multi-omic atlas of mouse early organogenesis by simultaneously profiling RNA expression and chromatin accessibility between E7.5 and E8.75, spanning late gastrulation and early organogenesis. Taking advantage of the simultaneous profiling of TF expression and cognate motif accessibility, we developed a tool to quantitatively predict TF binding events in cis-regulatory elements, which we used to quantify celltype-specific TF activities and infer gene regulatory networks that underlie cell fate transitions. We show that computational models trained on unperturbed data can be used to predict the effect of transcription factor perturbations. We validate this experimentally by showing that Brachyury is essential for the differentiation of neuromesodermal progenitors to somitic mesoderm fate by priming cis-regulatory elements.

## Author contributions

R.A., T.L., S.J.C. and W.R. conceived the project

T.L. and D.D performed embryo dissections

T.L. and S.J.C performed nuclear extractions

F.K. processed and managed sequencing data

A.N. performed gene targeting

R.A. and L.V. conceived and implemented the *in silico* ChIP-seq method.

R.A. and S.J.C. performed pre-processing and quality control

R.A. and G.L. performed the computational analysis

R.A. generated the figures

R.A. and S.J.C. interpreted results and drafted the manuscript.

W.R., S.J.C. and L.V. supervised the project.

All authors read and approved the final manuscript.

## Code availability

Code to reproduce the analysis is available at https://github.com/rargelaguet/mouse_organogenesis_10x_multiome_publication

## Data availability

Raw sequencing data together is available in the Gene Expression Omnibus under accession GSE205117 (reviewer token token gzcxaugylriplkn). Links to processed objects as well as to an R Shiny app for interactive data analysis are available in the github repository

## Acknowledgments

We thank Paula Kokko-Gonzales, Nicole Forrester and Amelia Edwards of the Babraham Institute Sequencing Facility for assistance with 10× Genomics library preparation, Katarzyna Kania and members of the CRUK-CI Genomics Core for 10x Genomics library preparation and Illumina sequencing and the Babraham Biological Support Unit for animal work. We thank Bart Theeuwes and Brendan Terry for comments on the manuscript. We thank all members of the Reik lab for their discussions and support.

The following sources of funding are gratefully acknowledged. This work was supported by the Wellcome Trust (awards 210754/Z/18/Z and 220379/Z/20/Z) and the BBSRC (award BBS/E/B/000C0421). T.L. was funded by the Wellcome Trust 4-Year PhD Programme in Stem Cell Biology and Medicine and the University of Cambridge, UK (203813/Z/16/A and 203813/Z/16/Z). This research was funded in whole or in part by the Wellcome Trust. R.A. was supported by the Wellcome for a Collaborative Award in Science (award 220379/Z/20/Z). The funding sources mentioned above had no role in the study design, in the collection, analysis and interpretation of data, in the writing of the manuscript and in the decision to submit the manuscript for publication.

## Conflict of interest statement

W.R. is a consultant and shareholder of Cambridge Epigenetix. R.A., D.D., F.K., S.J.C. and W.R. are employees of Altos Labs. The remaining authors declare no competing financial interests.

## Methods

### RNA data processing

Raw sequencing files were processed with CellRanger arc 2.0.0 using default arguments. Reads were mapped to the mm10-2020-A-2.0.0 genome and counted with GRCm38.92 annotation. Low-quality cells were filtered based on the distribution of QC metrics. Cells were required to have a minimum of 2000 UMIs per cell, a maximum of 40% mitochondrial reads and a maximum of 20% ribosomal reads. The resulting count matrix was stored using a SingleCellExperiment (Amezquita et al., 2019) (v 1.14.1) object. Normalisation and log transformation was performed using scran (Lun et al., 2016) (v1.20.1) and scuttle (McCarthy et al., 2017)(v1.2.1). Doublet detection was performed using the hybrid approach in the scds (v1.8.0) package.

### ATAC data processing

We used the ArchR package (Granja et al., 2021)(v1.0.1) for preprocessing of ATAC data. Briefly, arrow files were created from the ATAC fragment files. Cells were required to have a minimum of 3500 fragments per cell, a minimum TSS enrichment of 9, and a maximum blacklist ratio of 0.05. Pseudo-bulk replicates were obtained per cell type and peak calling was performed using macs2 (Zhang et al., 2008) (v2.2.7.1) using the cell type identified from the RNA expression as a group. A consensus peak set was obtained by an iterative overlapping strategy which is better at preserving cell type-specific peaks. Motif annotations were extracted from the CISBP (Weirauch et al., 2014) (v2) and JASPAR 2000 database (Castro-Mondragon et al., 2021). Motif matches for each peak were obtained using motifmatchr (v1.14.1), with a minimum motif width of 7 and a maximum q-value of 1e-4. Bigwig files were exported for each cell type for visualisation on the IGV browser (Robinson et al., 2011) (v2.11.0).

### Velocity analysis

Spliced and unspliced count matrices were extracted using velocyto (La Manno et al., 2018)(v0.17.17). Velocity analysis was performed using scVelo (Bergen et al., 2020) (v0.2.1) in dynamical mode.

### Metacell inference

When exploring continuous trajectories we summarised the data into metacells with the goal of achieving a resolution that retains the heterogeneity while overcoming the sparsity issues of single-cell data. We identified metacells (i.e. groups of cells that represent singular cell-states from single-cell data) using SEACells (Persad et al., 2022). Following the method guidelines, metacells were computed separately for each sample using approximately one metacell for every seventy-five cells. Following metacell identification, we regenerated gene expression and chromatin accessibility count matrices summarised at the metacell level.

Sample-specific count matrices were then concatenated and normalised using log-transformed counts per million.

### Multi-modal prediction of TF binding sites with *in silico* ChIP-seq

*The in silico* ChIP-seq library is a computational approach to link TFs to cis-regulatory elements in the form of ATAC peaks. Intuitively, we consider an ATAC peak *i* to be a putative binding site for TF *j* if *i* contains the j motif and its chromatin accessibility correlates with the RNA expression of *j*. Formally, we calculate the *in silico* TF binding score for ATAC peak *i* and TF *j* with the following equation:

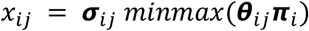

where ***σ**_ij_* is the correlation between the chromatin accessibility of peak *i* and the RNA expression of TF *j*. ***θ**_ij_* is the motif score for peak *i* and TF *j*, and ***π**_i_* is the maximum chromatin accessibility of peak *i* (across all samples). Note that the TF binding score ranges from -1 to 1 due to the *minmax* normalisation. A negative *in silico* TF binding score value denotes a repressive event, where the chromatin accessibility of ATAC peak *i* is negatively regulated by TF *j*. In contrast, a positive value denotes an activatory event, where the chromatin accessibility of peak *i* is positively regulated by TF *j*. Although the TF *in silico* score is continuous, some analysis require a binarised association between TFs and cis-regulatory elements. In this case the *in silico* TF binding score can be modulated as a hyperparameter, such that small values will lead to many predicted TF binding events, a high false positive rate and a low true positive rate. Large values will lead to fewer predicted TF binding events, but a low false positive rate and a high true positive rate. We performed grid search and found that values between 0.10 and 0.30 provide reasonable trade-offs between the number of predicted TF binding events and the accuracy of the predictions.

### Quantification of transcription factor activities per cell type using chromVAR and chromVAR-Multiome

TF activities were calculated using the chromVAR algorithm (Schep et al., 2017). The method takes as input the ATAC peak matrix and a set of position-specific weight matrices (PWMs) encoding TF sequence affinities. Here we used the JASPAR (2022)(Castro-Mondragon et al., 2021) and CISBP (v2.0)(Weirauch et al., 2014) databases. Briefly, for each TF motif contained within an ATAC peak and each cell (or cell type, when calculated at the pseudo-bulk level), chromVAR calculates a z-score that measures the difference between the total number of fragments that map to motif-containing peaks and the expected number of fragments (based on the average of all cells). Importantly, the normalisation and scaling that chromVAR applies is aimed at mitigating technical biases between cells (Tn5 tagmentation efficiency, PCR amplification, etc.) and features (GC content, mean accessibility, etc.). Here we modified the chromVAR algorithm to use the putative TF binding sites from the *in silico* ChIP-seq library, instead of all TF motif instances. We refer to this approach as chromVAR-Multiome

### Multi-modal dimensionality reduction

We generated a multi-modal latent embedding using MOFA+ (Argelaguet et al., 2020). Briefly, the method takes as input multiple data modalities and performs multi-view matrix factorisation to generate a set of latent factors that can be used for a variety of downstream tasks. Here we used as input to MOFA the RNA expression and ATAC peak matrix. Feature selection was performed to enrich for highly variable features (3,000 genes and 25,000 ATAC peaks). Optionally, one can also use as input latent variables that result from linear dimensionality reduction (Principal Component Analysis in the case of the RNA expression and Latent Semantic Indexing in the case of ATAC peaks). This leads to a significant increase in speed and also mitigates challenges linked to class imbalance (i.e. the two views having many different features). We ran MOFA with a fixed set of 30 factors, which we subsequently used as input to the UMAP algorithm(McInnes et al., 2018) to generate a (non-linear) two-dimensional embedding that is suitable for visualisation.

### TF marker scores

We used the chromVAR-Multiome values to define TF marker scores for each combination of cell type and TF. We adopted a similar algorithm as used for the definition of marker genes in Seurat (*FindMarkers* function) and scran (*findMarkers* function). First, we performed differential analysis between each pair of cell types using a t-test. Then, for each TF *i* and cell type *j* we counted the number of significant differential comparisons between cell type *j* and all other cell types different from *j*. Instead of aggregating the p-values and fold changes, as done in Seurat and scran, we adopt a more intuitive metric and define the TF marker score as the fraction of differential comparisons. Intuitively, the higher the score of TF *i* in cell type *j* the more active that TF *i* is in cell type *j* when comparing the chromVAR-Multiome values to the other cell types. The maximum TF marker score value is 1, when all differential comparisons are significant. When defining the catalogue of TF activities per cell type (Figure 3f), we set a minimum TF marker score of 0.75.

### Gene accessibility scores

Here we quantified promoter accessibility by adding all reads that map to the region that is 500bp upstream and 100bp downstream of the transcription start site (TSS). TSS annotations are obtained from the BioMart database using the Bioconductor GenomicFeatures package (v1.48.1). Note that here we disabled ArchR’s default gene accessibility model, which incorporates information from cis-regulatory elements that are located near the TSS. Although this approach is more predictive of changes in gene expression, it is problematic when applied to genomic regions with high gene density, as cis-regulatory elements cannot be confidently linked to genes.

### Pooling cells from the same cell type into pseudo-bulk replicates

The sparsity of the single-cell data limits the statistical analysis, the visualisation strategies and overall the biological insights that can be extracted from the data (Squair et al., 2021). For some analysis that involve cell type comparisons (including differential analysis, peak calling or *in silico* ChIP-seq inference), we create “pseudo-bulk” replicates by aggregating reads from all cells that belong to the same cell type. The pseudo-bulk strategy is particularly important for snATAC-seq data, as ATAC peaks typically have very few reads per cell. For differential analysis between cell types, we follow the approach suggested in(Crowell et al., 2020) and create the same number of replicates per cell type by bootstrapping cells assigned to the same cell type. Besides reducing sparsity, this approach also helps address the problem of having a different number of samples per group when doing differential analysis at single-cell resolution, which often leads to p-values being systematically different depending on the number of samples per group.

### Genome Browser visualisation

We use the getGroupBW function in ArchR to group, summarise and export a bigwig file for each cell type. Briefly, the function calculates normalised accessibility values along the genome using 100bp tiles. We visualise the ATAC bigwig files as separate tracks in the IGV Browser (v2.11.0)(Thorvaldsdottir et al., 2013)

### Differential RNA expression and chromatin accessibility

Following the guidelines from previous studies (Squair et al., 2021), we performed differential analysis using pseudo-bulk replicates for each cell type (and genotype, in the Brachyury KO study). For each group we derived 5 replicates by bootstrapping different subsets of cells at random. Each pseudo-bulk replicate contained 30% of the total number cells, with at least 25 cells per replicate. Subsequently, read counts were aggregated for each group, followed by normalisation with log-transformed counts per million (CPMs). Note that this “pseudo-bulk-with-replicates” approach yields the same number of samples per group, which facilitates differential analysis comparisons. Differential analysis was performed using the negative binomial model with a quasi-likelihood test implemented in edgeR (Robinson et al., 2010). Significant hits were called with a 1% FDR (Benjamini–Hochberg procedure) and a minimum log2 fold change of 1. Hits with small average expression values (log normalised counts <=2) were ignored, as this can lead to artificially large fold change values.

### Identification of celltype marker genes and regulatory elements

Cell type-specific marker genes and peaks were identified using the reference cells (i.e. the cells form the Brachyury KO study were excluded). First, we performed differential analysis between each pair of cell types using the strategy outlined above. Then, for each cell type, we labelled as marker genes or as marker peaks those hits that are differentially expressed/accessible and upregulated in the cell type of interest in more than 85% of the comparisons.

### Mapping to a reference atlas and cell type assignment

Cell types were assigned by mapping the RNA expression profiles to a reference atlas from the same stages (Pijuan-Sala et al., 2019). The mapping was performed by matching mutual nearest neighbours with the fastMNN algorithm (*batchelor* R package v1.8.1)(Haghverdi et al., 2018). First, count matrices from both experiments were concatenated and normalised together using scran (v1.20.1). Highly variable genes were selected(Lun et al., 2016) from the resulting expression matrix and were used as input for Principal Component Analysis. A first round of batch correction was applied within the atlas cells to remove technical variability between samples. A second round of batch correction was applied to integrate query and atlas cells within a joint PCA space. Then, for each query cell we used the *queryKNN* function in BiocNeighbors to identify the 25 nearest neighbours from the atlas. Finally, a cell type was inferred for each query cell by majority voting among the atlas neighbour cells.

Mapping to the spatial atlas and imputation of spatially-resolved ChromVAR-Multiome scores Mapping of the 10x Multiome cells to the spatially-resolved transcriptomic atlas was done using the same approach described above for the scRNA-seq reference atlas. This integration is however more challenging due to the sparsity of the seqFISH data set and the different nature of the size factors. Here we followed the strategy outlined in(Lohoff et al., 2022) and applied cosine normalisation on the log-normalised counts. For simplicity, we used as reference a single z-slice from a representative E8.5 embryo.

Finally, we used the mapping to impute spatially-resolved TF activities. We transferred the chromVAR-Multiome scores from the 10× Multiome cells onto the nearest neighbours of spatial atlas. Due to the noisy estimates in single-cell data and the presence of outliers, we performed kNN denoising before visualisation.

### Inference of the TF regulatory network underlying differentiation of Neuromesodermal progenitors

First, we selected metacells of the NMP differentiation trajectory. Note that we discourage the use of data at single cell resolution, as the sparsity of snATAC-seq makes it challenging to obtain reliable associations between the RNA expression of TFs (which are typically lowly expressed genes) and chromatin accessibility of cis-regulatory regions. Second, we used the *in silico* ChIP-seq methodology to link TFs with cis-regulatory elements. Third, we linked cis-regulatory regions to nearby genes via a maximum genomic distance of 50kb. Note that this step results in a many-to-many mapping, where each gene can be linked to multiple cis-regulatory regions, and each cis-regulatory region can be linked to many genes. Fourth, we built a linear regression model of target gene RNA expression as a function of the TF’s RNA expression. Finally, we visualise the GRN as a directed graph where nodes correspond to TFs and target genes (which can also be other TFs), where the edge width is given by the slope of the linear regression models.

### *In silico* TF perturbation with CellOracle

Briefly, CellOracle leverages a gene regulatory network and a differentiation trajectory to predict shifts in cellular identities by simulating the effects of TF perturbations on the GRN configuration. It simulates gene expression values upon TF perturbation, which are then compared with the gene expression of local neighbourhoods to estimate transition probabilities between cell states. Finally, CellOracle creates a transition trajectory graph to project the predicted identity of these cells upon TF perturbation. Here we used the GRN inferred from the NMP differentiation trajectory as input, where target genes are constrained to also be TFs. Given the improved signal-to-noise ratio in the metacell representation, we disable the default kNN denoising step.

### Embryos and nuclear isolation

C57BL/6Babr mice were bred and maintained by the Babraham Institute Biological Support Unit. All mouse experimentation was approved by the Babraham Institute Animal Welfare and Ethical Review Body. Animal husbandry and experimentation complied with existing European Union and United Kingdom Home Office legislation and local standards.

Following dissection, embryos from the same stages were pooled to give sufficient cell numbers. Embryos were dissociated into single-cells using 200μl of TriplE Express for 10 minutes at 37°C on a shaking incubator. 1ml of ice-cold 10% FBS in PBS was added to quench and cells were filtered using a 40μM Flowmi cell strainer. Following centrifugation at 300g for 5 minutes, the supernatant was discarded and cells were resuspended in 50μl of PBS containing 0.04% BSA. Cells were counted and viability assessed using trypan blue staining on a Countess II instrument (Invitrogen). >95% of cells were negative for trypan blue indicating high sample quality.

Nuclear isolation was carried out according to the low-cell input version of the 10X protocol for cell lines and PBMCs (https://assets.ctfassets.net/an68im79xiti/6t5iwATCRaHB4VWOJm2Vgc/bdfd23cdc1d0a321487c8b231a448103/CG000365_DemonstratedProtocol_NucleiIsolation_ATAC_GEX_Sequencing_RevB.pdf). Specifically, the 50μl cell suspension was transferred to a 0.2ml PCR tube and centrifuged at 300g for 5 minutes. After removing the supernatant, cells were resuspended in 50μl ice cold nuclear extraction (NE) buffer (10mM Tris pH 7.5, 10mM NaCl, 3mM MgCl2, 1% BSA, 0.1% Tween, 1mM DTT, 1U/ul RNaseIn (Promega), 0.1% NP40, 0.01% Digitonin) and incubated on ice for 4 minutes. 50μl of wash buffer (identical to NE buffer but lacking NP40 and digitonin) was added and nuclei were centrifuged at 500g for 5 minutes at 4°C. After removing the supernatant, nuclei were washed once in 50μl of diluted nuclei buffer (10× Genomics), span down and finally resuspended in 7ul of dilute nuclei buffer (10× Genomics). 1μl was used to assess quality using a microscope and count nuclei using a Countess II instrument. >99% of nuclei stained positive for trypan blue and the nuclei were found to have the expected morphology. Nuclei were diluted such that a maximum of 16,000 were taken forward for 10× Multiome library preparation.

### Brachyury gene targeting

One-cell stage zygotes were obtained from C57BL/6Babr superovulated matings. CRISPR/Cas9 reagent consisted of Cas9 protein (200ng/ul) and a sgRNA targeting exon 3 of the Brachyury gene (120ng/ul, ACTCTCACGATGTGAATCCG), diluted in Opti-MEM I (Thermo Fisher). Control embryos received Cas9 but no gRNA. Super electroporator NEPA21 and platinum plate electrodes 1mm gap (CUY501P1-1.5) were used for electroporation. Four repeats of poring pulses (40V, 3.5ms length and 50ms intervals) and five repeats of transfer pulses (7V, 50ms length, 50ms intervals) were applied to zygotes. Approximately 50 embryos were added to 5-6ul of CRISPR/Cas9mix per electroporation. Embryos were cultured overnight and only 2-cell stage embryos were transferred into pseudo-pregnant recipients, which were later harvested to obtain E8.5 embryos. In total this yielded 3 control embryos and 7 Brachyury KO embryos which were pooled for processing.

For genotyping embryonic yolk sacs were lysed using QuickExtract buffer prior to PCR amplification of a region spanning the predicted cut site (forward: GTAGGCAGTCACAGCTATGA, reverse: GGGTTTAATGGTGTATAGCG). The resulting amplicon was Sanger sequenced and the trace was analysed using Synthego ICE analysis producing a KO score of 93% (https://www.synthego.com/products/bioinformatics/crispr-analysis)(Conant et al., 2022).

### 10× Multiome library preparation and sequencing

Libraries were prepared using the 10x Genomics Chromium and sequenced on a Novaseq 6000 instrument (Illumina) using the recommended read-lengths. This yielded medians of 720 million RNA-seq reads and 481 million ATAC reads per sample. We recovered a median of 7,700 cells per sample prior to quality control.

### ChIP-seq data processing

ChIP-seq data for TFs Cdx2, Foxa2, Gata1, Gata4, Tal1 and Tbx5 was obtained from the Gene Expression Omnibus. Due to the limited availability of *in vivo* ChIP-seq datasets, we used *in vitro* models that more closely resemble the gastrulating embryo (**Supplementary Table 1**). Reads were trimmed using Trim Galore (v0.4.5) and mapped to *M. musculus* GRCm38 using Bowtie2 (Langmead and Salzberg, 2012) (v2.3.2). Bigwig files were generated for genome browser visualisation using samtools (v1.13)(Li et al., 2009) and bamCoverage (v3.5.1)(Ramírez et al., 2016). Peak calling was performed using macs2 (v2.2.7.1)(Zhang et al., 2008) with the “--broad and --broad-cutff 0.1” arguments.

## Notes

### Summary of Updates

Figures and manuscript have been updated to improve clarity

