## Supplementary Information for "Decoding gene regulation in the mouse embryo using single-cell multi-omics"

### *in silico* ChIP-seq: multi-modal prediction of transcription factor binding sites

#### Model description

*The in silico* ChIP-seq library is a computational approach to link TFs to cis-regulatory elements in the form of ATAC peaks. Intuitively, we consider an ATAC peak *i* to be a putative binding site for TF *j* if *i* contains the j motif and its chromatin accessibility correlates with the RNA expression of *j*. Formally, we calculate the *in silico* TF binding score for ATAC peak *i* and TF *j* with the following equation:

$$x_{ij} = \boldsymbol{\sigma}_{ij}\, minmax(\boldsymbol{\theta}_{ij}\boldsymbol{\pi}_i)$$

where $\boldsymbol{\sigma}_{ij}$ is the correlation between the chromatin accessibility of peak *i* and the RNA expression of TF *j*. $\boldsymbol{\theta}_{ij}$ is the motif score for peak *i* and TF *j*, and $\boldsymbol{\pi}_i$ is the maximum chromatin accessibility of peak *i* (across all samples). Note that the TF binding score ranges from -1 to 1 due to the *minmax* normalisation. A negative *in silico* TF binding score value denotes a repressive event, where the chromatin accessibility of ATAC peak *i* is negatively regulated by TF *j*. In contrast, a positive value denotes an activatory event, where the chromatin accessibility of peak *i* is positively regulated by TF *j*. Although the TF *in silico* score is continuous, some analysis require a binarised association between TFs and cis-regulatory elements. In this case the *in silico* TF binding score can be modulated as a hyperparameter, such that  small values will lead to many predicted TF binding events, a high false positive rate and a low true positive rate. Large values will lead to fewer predicted TF binding events, but a low false positive rate and a high true positive rate. We performed grid search and found that values between 0.10 and 0.30 provide reasonable trade-offs between the number of predicted TF binding events and the accuracy of the predictions.

#### Comparison with other approaches

- CellOracle (Kamimoto et al., 2020): the main purpose of this model is gene regulatory network inference to test *in silico* TF perturbations. TFs are associated to cis-regulatory regions using scATAC-seq data, with no contribution from the RNA expression. However, CellOracle provides flexibility in the definition of links between TFs and cis-regulatory regions. Here we used the *in silico* ChIP-seq results as input for the GRN construction step implemented in CellOracle.
- Virtual ChiP-seq (Karimzadeh and Hoffman, 2019): it is a supervised deep learning framework which uses publicly available ChIP-seq experiments in multiple cell types, genomic conservation, and (bulk) RNA expression to predict TF binding. In contrast to our unsupervised *in silico* ChIP-seq method, this requires ChIP-seq data as input.
- GRaNIE (Kamal et al., 2021): similar to our *in silico* ChIP-seq, this method infers links between TFs and ATAC peaks based on co-variation between TF expression and chromatin accessibility of putative transcription factor binding sites. The main difference with the *in silico* ChIP-seq is that this GRaNIE only uses the links between TFs and ATAC peaks as a proxy for inference of gene regulatory networks. In our

method we incorporated the motif score, the chromatin accessibility values and the co-variation between TF expression and chromatin accessibility to devise an interpretable TF binding score, which we validated by benchmarking using ChIP-seq data.

#### Celltype specificity

The output of the *in silico* ChIP-seq model is a matrix of TF-peak binding scores. The key element of the model is the correlation between the TF's RNA expression and the chromatin accessibility of its putative target peaks. This correlation is calculated across all metacells or pseudobulk cell types (depending on the chosen data resolution, see discussion below). By default all cell types are used to create the model, and thus there is no cell type specificity output by the model. However, there are strategies to obtain cell type specificity:

(1) Subset the input data: the *in silico* ChIP-seq library can be inferred using selected subsets of cell types. For example, if one desires to obtain predicted binding sites for specific cell types or trajectories (as we have done in Figure 5 of the manuscript for neuromesodermal progenitors), one can run the model using the metacells that have been assigned to these cell types or trajectories. Importantly, this approach requires biological variability to exist across metacells in order to calculate meaningful correlations between RNA expression and chromatin accessibility. Consequently, this approach would not work for homogeneous cellular populations.

(2) Assume that the cell type where the TF's expression and TF motif accessibility is highest is the cell type where the TF is active. With this assumption, one can then use the chromVAR-Multiome approach (described in Methods) to quantify TF activities for each cell, metacell or cell type (depending on the chosen data resolution, see discussion below).

#### Benchmark

To benchmark the *in silico* ChIP-seq library, we used publicly available ChIP-seq experiments for a set of TFs that are known to play key roles during mouse gastrulation and early organogenesis. Due to the limited availability of *in vivo* ChIP-seq datasets, we mostly relied on data from *in vitro* models that resemble the gastrulating embryo. For each TF experimental ChIP-seq data we performed peak calling with *macs2* (Zhang et al., 2008), with biological replicates when available. We then selected the top 1,000 peaks with lowest q-value for each experiment. These sites represent the strongest TF binding events and were defined as true binding sites in the benchmark.

We performed two benchmarks to compare the model with and without RNA expression:

- Assessment of the consistency between experimental ChIP-seq values and *in silico* TF binding scores (Figure 2d in the manuscript): for each predicted binding site, we plotted the *in silico* binding score and the experimental ChIP-seq score. For all TFs benchmarked, an increase in the *in silico* TF binding score is consistent with changes in the experimental ChIP-seq values in both models, but substantially better scores are achieved with the model that incorporates RNA expression information.
- A receiver operating characteristic (ROC) curve after binarisation of the predicted binding sites (Figure 2e in the manuscript): first, we binarised the *in silico* TF binding predictions using a range of score thresholds, resulting in predicted binding sites

(score>=threshold) and non-binding sites (score<threshold). Subsequently, we overlapped these sites with the experimental ChIP-seq peaks using their genomic coordinates, and generated a contingency table as shown below. Finally, we plotted the true positive rate (TPR) against the false positive rate (FPR) at various threshold settings.

| | **Observed binding site** | **Observed non-binding site** |
|---|---|---|
| **Predicted binding site** | True positive | False negative |
| **Predicted non-binding site** | False positive | True negative |

#### Selecting a data resolution

The key computation of the *in silico* ChIP-seq model is the correlation between the TF's RNA expression and the chromatin accessibility of its putative target peaks. This correlation can be calculated using different data representations at varying resolutions: cells, metacells, or pseudo-bulk samples at the cell type level. The cellular representation has the advantage of not requiring data aggregation and preserves all biological variability. However, the quantification of TF expression and chromatin accessibility at cis-regulatory elements is unreliable due to data sparsity and technical noise. In addition, a major disadvantage is the imbalance between cell types such that more abundant cell types contribute more to the correlation estimates. In contrast, the pseudobulk representation provides robust RNA expression and chromatin accessibility estimates, and also has the advantage of providing the same number of data points for each cell type, thus avoiding biases linked to differences in cellular abundance. Finally, the metacell representation is a tradeoff between the two representations. As shown in multiple studies (Ben-Kiki et al., 2021; Bilous et al., 2022; Persad et al., 2022), metacells provide an excellent tradeoff of retaining cellular heterogeneity while overcoming the sparsity issues of single-cell data.

To illustrate the tradeoffs of the three data representations, we plotted the distribution of correlation coefficients between TF expression and target gene expression, quantified at the cell (left), metacell (middle) or pseudobulk (right) resolution.

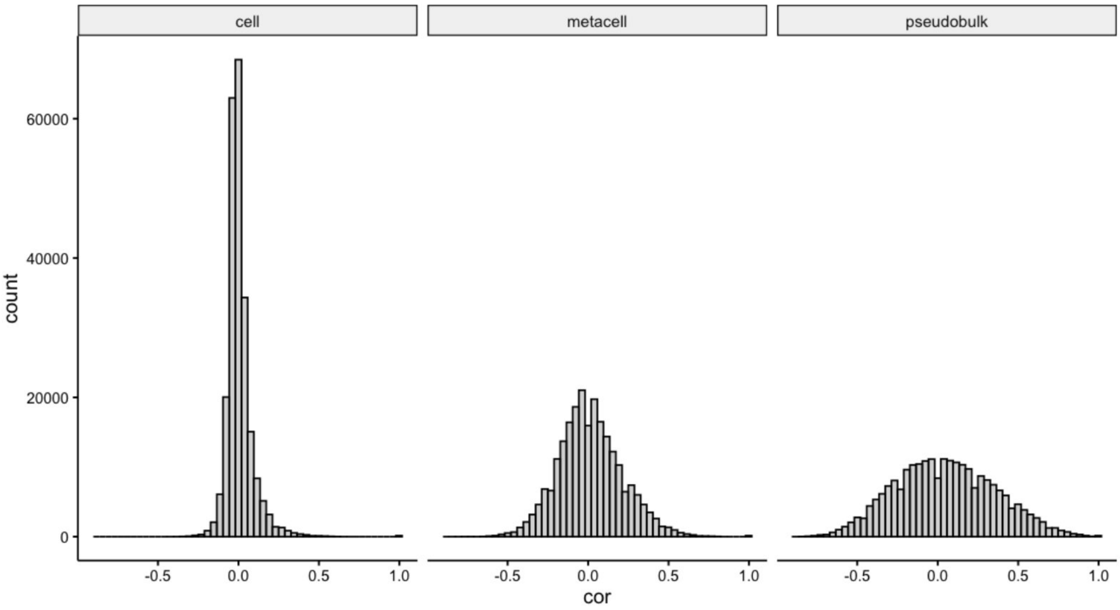


To showcase a specific example, the following scatterplots show the RNA expression of the TF Gata1 versus the expression of a target gene, quantified again at the three levels of data resolution.

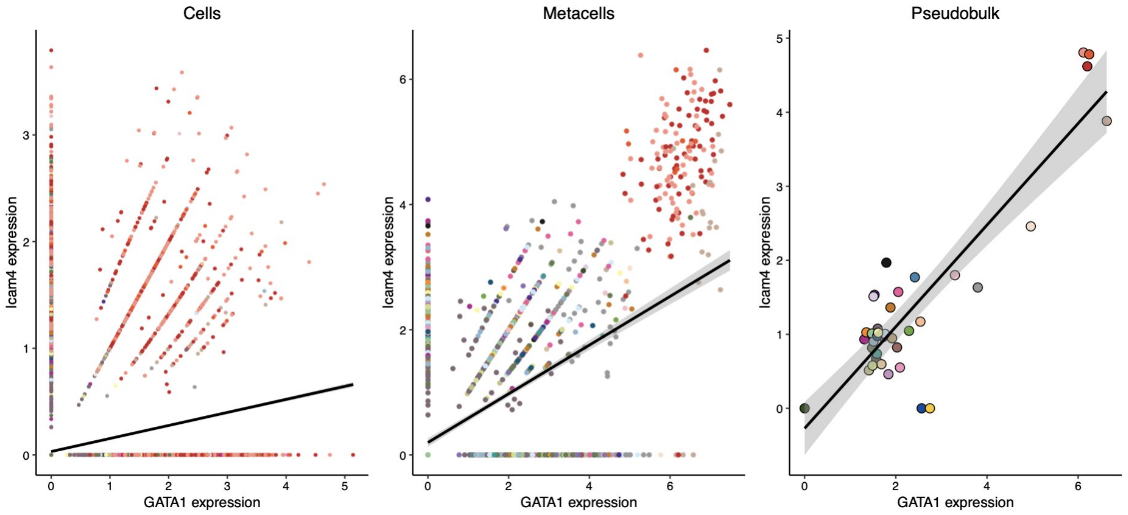


#### Limitations of the method

Although in silico ChIP-seq provides a conceptual advantage for the analysis of single-cell multi-modal data, it is not free of limitations:

- **Simplistic relationships between RNA expression and chromatin accessibility**: the model exploits linear correlations between TF RNA expression and chromatin accessibility. There are however cases of regulatory relationships where a time lag exists between the two modalities, most notably in the case of epigenetic priming events. Although the *in silico* ChIP-seq can capture some priming events (see for example Figure 7f in the manuscript), many of these non-linear relationships will be missed.

- **Fixed representations of TF motifs via position weight matrices (PWMs)**: sequence-based PWMs models are simple and intuitive. However, variation in sequence specificity and cooperative TF binding across cell types prevents PWM-based methods from accurately capturing true TF binding. More recent methods that combine PWMs from popular TF motif databases with *de novo* inference of TF motifs (de Almeida et al., 2022; Shrikumar et al., 2018; Yuan and Kelley, 2022; Zhang et al., 2022) have the potential to capture novel and more complex relationships, including TF cooperativity events. Future research could explore variations of the *in silico* ChIP-seq where deep learning can replace the simple equations that we formulated in this work.

- **TF cooperativity is ignored**: cis-regulatory regions typically contain multiple TF motifs that serve as potential TF binding sites. In many cases, TF interactions have been shown to alter the sequence recognition (Ibarra et al., 2020; Slattery et al., 2014). However, studying TF cooperativity remains extremely challenging in the absence of good-quality experimental TF binding data. Thus, here we applied the *in silico* ChIP-seq methodology independently for each TF and thus have not considered cooperative binding. In the future, we hope to expand the model with novel multi-modal technologies where TF binding is simultaneously profiled with RNA expression, chromatin accessibility and other readouts at single-cell resolution(Gopalan et al., 2021; Zhu et al., 2021).
