## Supplementary Figures for "Decoding gene regulation in the mouse embryo using single-cell multi-omics"

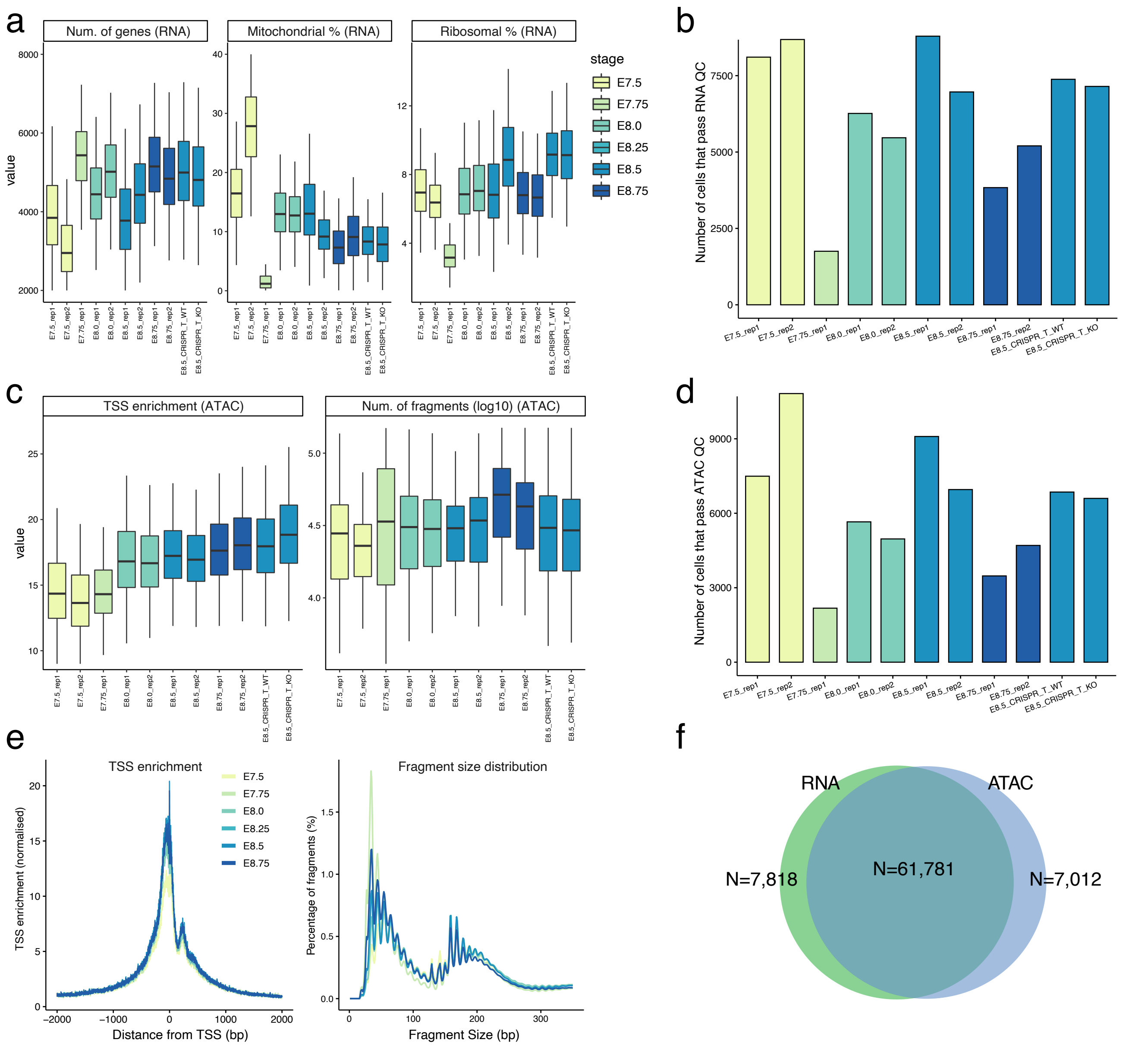


**Figure S1: Quality control statistics per sample.**

(a) Boxplots displaying RNA-seq quality control (QC) metrics per cell: the number of expressed genes (left), the percentage of mitochondrial reads (middle) and the percentage of Ribosomal reads (right). Each box is a sample, coloured by embryonic stage.

(b) Barplots displaying the number of cells that pass RNA-seq QC for each sample. Bars are coloured by embryonic stage.

(c) Boxplots displaying ATAC-seq quality metrics per cell: the enrichment of reads in the Transcription Start Site (TSS) (Granja et al., 2021) (left) and the number of fragments (right). Each box is a sample, coloured by embryonic stage.

(d) Barplots displaying the number of cells that pass ATAC-seq QC for each sample. Bars are coloured by embryonic stage.

(e) Histograms of QC statistics for ATAC-seq per sample. The left plot shows the number of fragments as a function of the distance from the nearest gene's TSS. The right plot shows the insert size distribution of ATAC-seq fragments.

(f) Venn diagram showing the overlap between cells that pass QC for the two modalities.

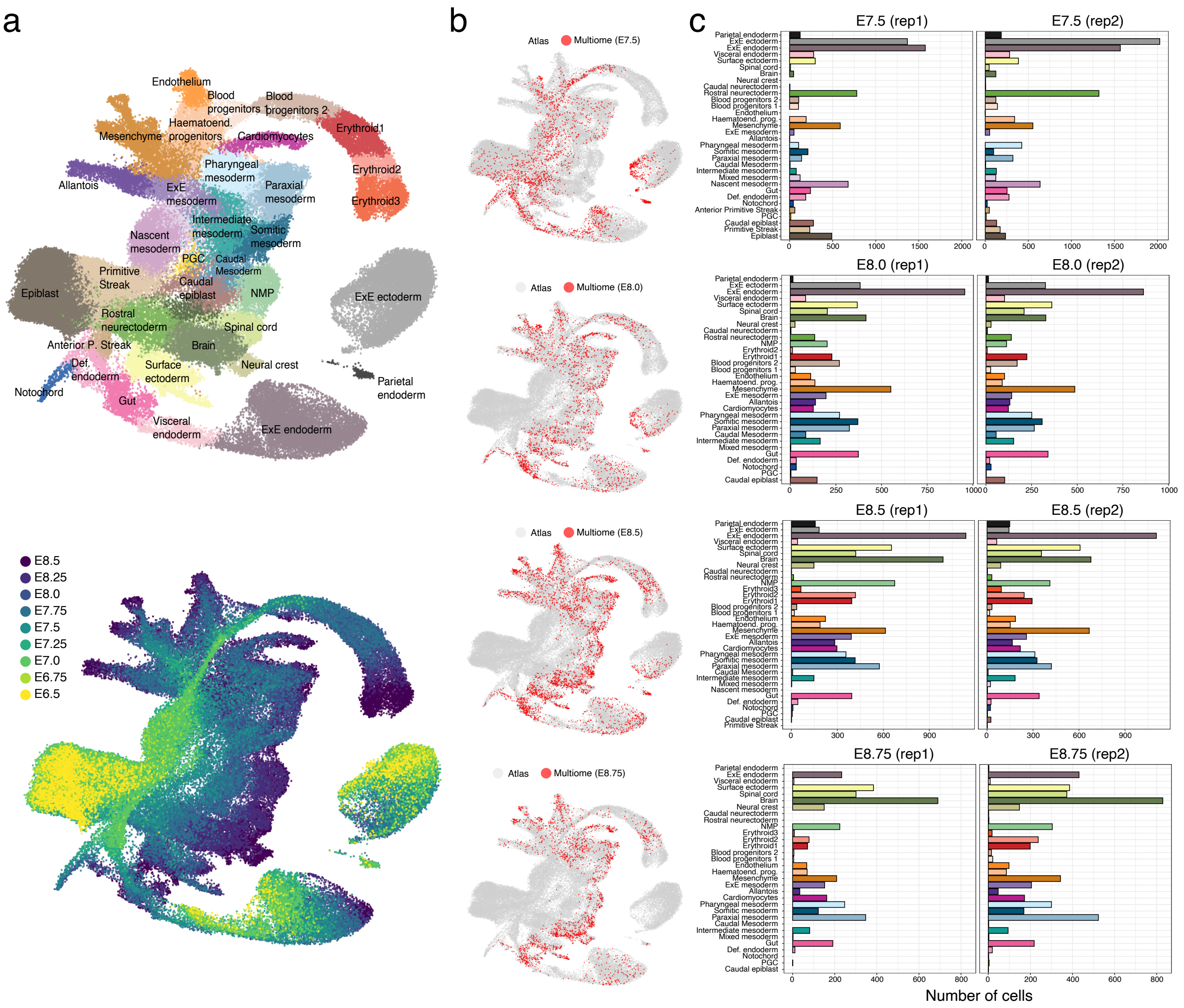


**Figure S2: Mapping to the reference atlas and cell type annotation.**

(a) UMAP plot from the reference atlas (Pijuan-Sala et al., 2019). Dots are coloured by cell type (top) or embryonic stage (bottom).

(b) Same UMAP plot as in (a). Coloured in red are cells that represent matching nearest neighbours to cells from this study (Methods). f corresponds to different stages.

(c) Bar plots displaying the number of cells for each cell type and sample. Each row corresponds to different stages.

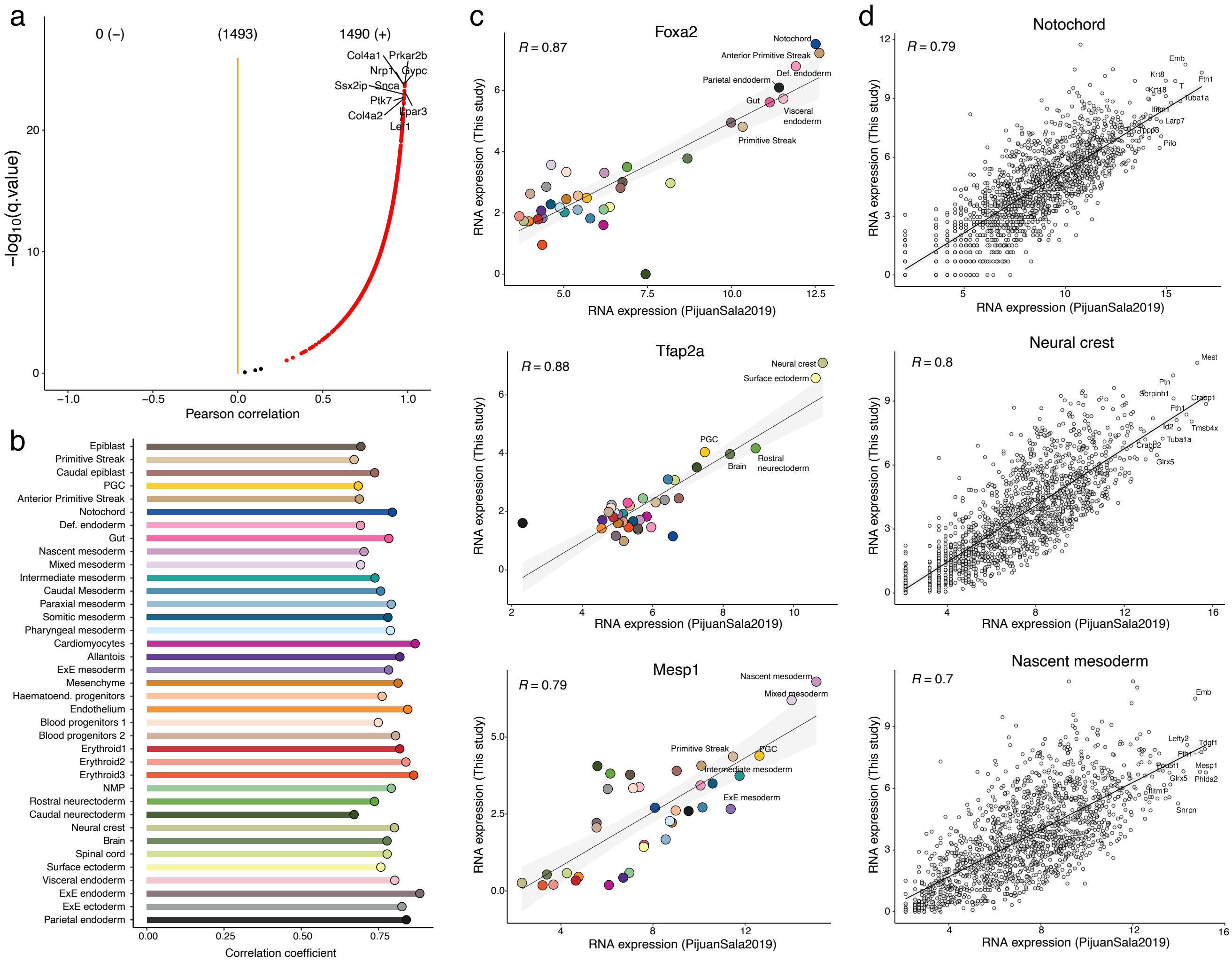


**Figure S3: Comparison of the snRNA expression profiles from the 10x Multiome with existing scRNA-seq data from overlapping stages.**

(a) Volcano plot displays the results of correlation tests per gene (across cell types) between the reference dataset (Pijuan-Sala et al., 2019) and this study. Correlations were computed at the cell type level after pseudobulk (i.e. each observation corresponds to a different cell type). Only cell type marker genes (N=1493) were considered for this analysis.

(b) Bar plots display the results of correlation tests per cell type (across genes) between the reference data set (Pijuan-Sala et al., 2019) and this study. As in (a), marker genes were considered for this analysis.

(c) Scatter plots show the RNA expression levels for three representative genes between the reference dataset (x-axis) and this study (y-axis). Each dot corresponds to a different cell type. Line represents the linear regression fit. Shown in the top left corner is the Pearson correlation coefficient.

(d) Scatter plots show the RNA expression levels for three representative cell types between the reference dataset (x-axis) and this study (y-axis). Each dot corresponds to a different gene. Line represents the linear regression fit. Shown in the top left corner is the Pearson correlation coefficient.

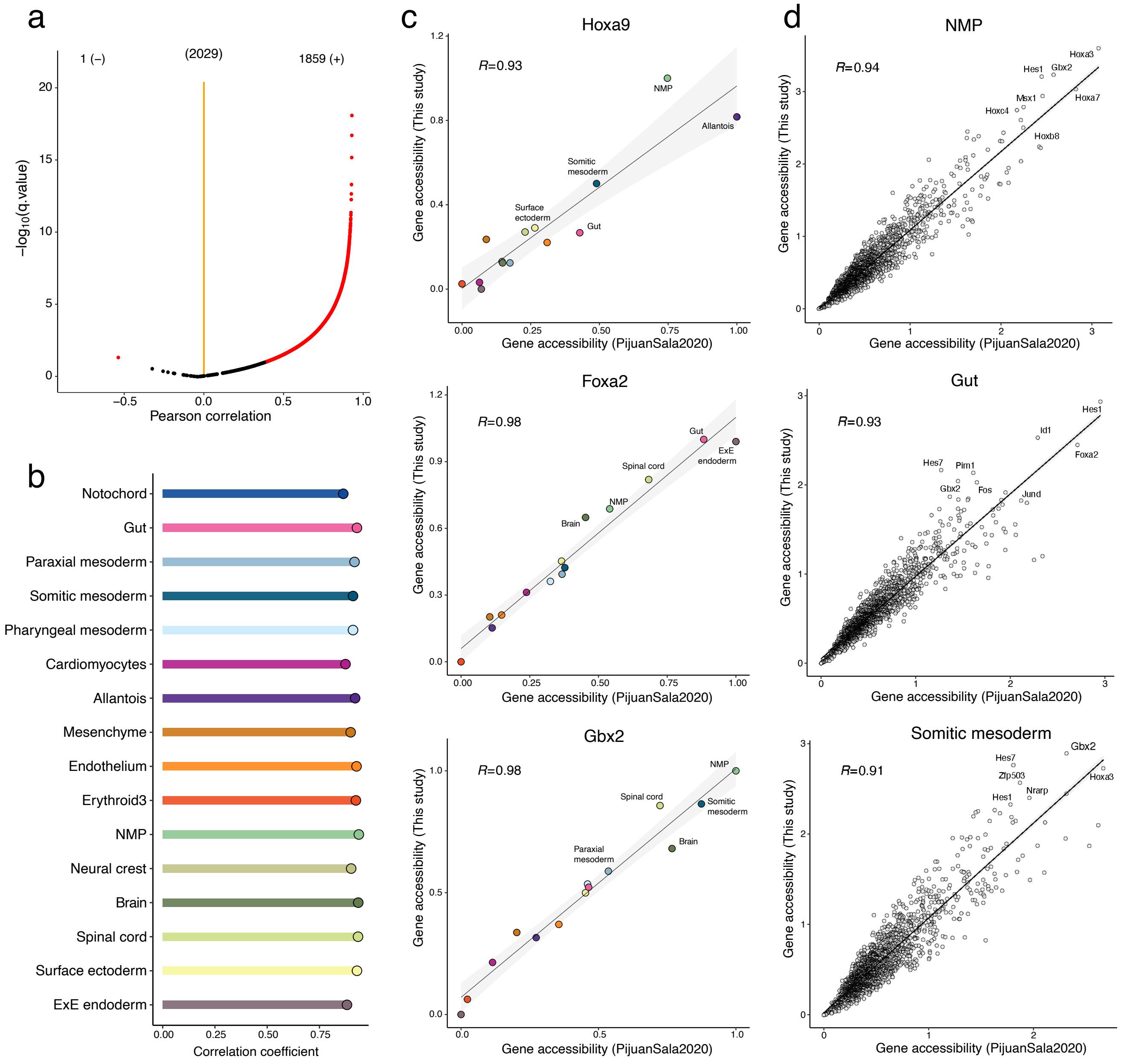


**Figure S4: Comparison of the 10x Multiome chromatin accessibility data with existing scATAC-seq data from E8.25.**

(a) Volcano plot displays the results of correlation tests per gene (across cell types) between the reference dataset (Pijuan-Sala et al., 2020) and this study. Correlations were computed at the cell type level after pseudobulk (i.e. each observation corresponds to a different cell type). Chromatin accessibility gene scores for marker genes were considered for this analysis (Methods).

(b) Bar plots display the results of correlation tests per cell type (across genes) between the reference data set (Pijuan-Sala et al., 2020) and this study. As in (a), marker genes were considered for this analysis.

(c) Scatter plots show the chromatin accessibility levels for three representative genes between the reference dataset (x-axis) and this study (y-axis). Each dot corresponds to a different cell type. Line represents the linear regression fit. Shown in the top left corner is the Pearson correlation coefficient.

(d) Scatter plots show the chromatin accessibility levels for three representative cell types between the reference dataset (x-axis) and this study (y-axis). Each dot corresponds to a different gene. Line represents the linear regression fit. Shown in the top left corner is the Pearson correlation coefficient.

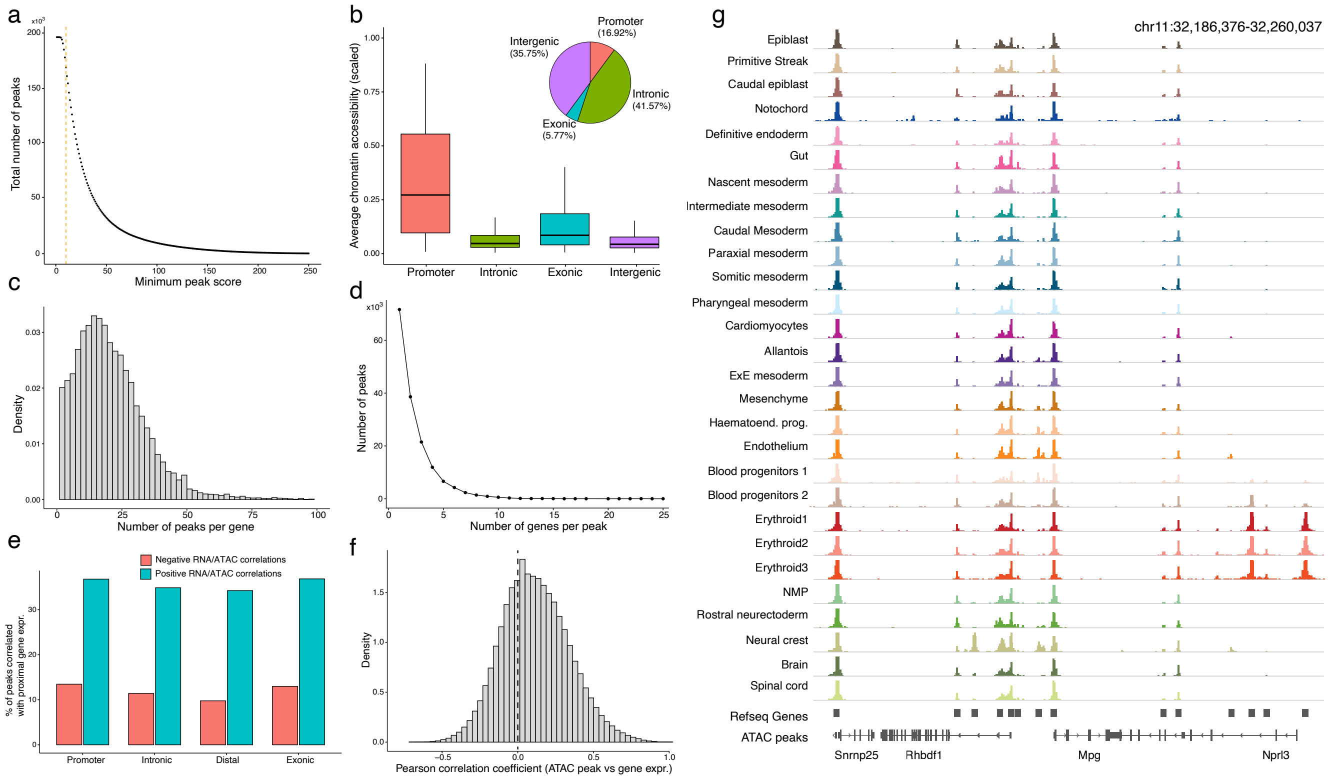


**Figure S5: Identification of cis-regulatory elements using celltype-aware peak calling.**

(a) Scatter plot showing the relationship between the ATAC peak score cutoff (x-axis) and the corresponding number of ATAC peak calls (y-axis). Dashed line indicates the cutoff used in subsequent analyses.

(b) Boxplots showing the mean chromatin accessibility across all cells for peaks overlapping different genomic contexts. Inset: pie chart showing the percentage of peaks overlapping each genomic context.

(c) Histogram showing the number of ATAC peaks linked to each gene (maximum genomic distance of 50kb).

(d) Line plot showing the the number of genes linked to each peak.

(e) Barplot showing the percentage of ATAC peaks whose accessibility correlates with expression of at least one linked gene (q-value¡0.01 and a minimum absolute correlation of 0.25). Positive correlates are coloured in blue whereas negative correlations are coloured in red.

(f) Histogram displaying the distribution of Pearson correlation coefficients between ATAC peak accessibility and RNA expression (quantified at the pseudobulk level across cell types).

(g) Genome browser plot of a representative genomic locus that contains ATAC peaks that display variability in chromatin accessibility across cell types as well as peaks that are relatively homogeneous across cell types. Note that highly variable ATAC typically map to intergenic or intronic regions, whereas homogeneous ATAC peaks are found in promoter regions.

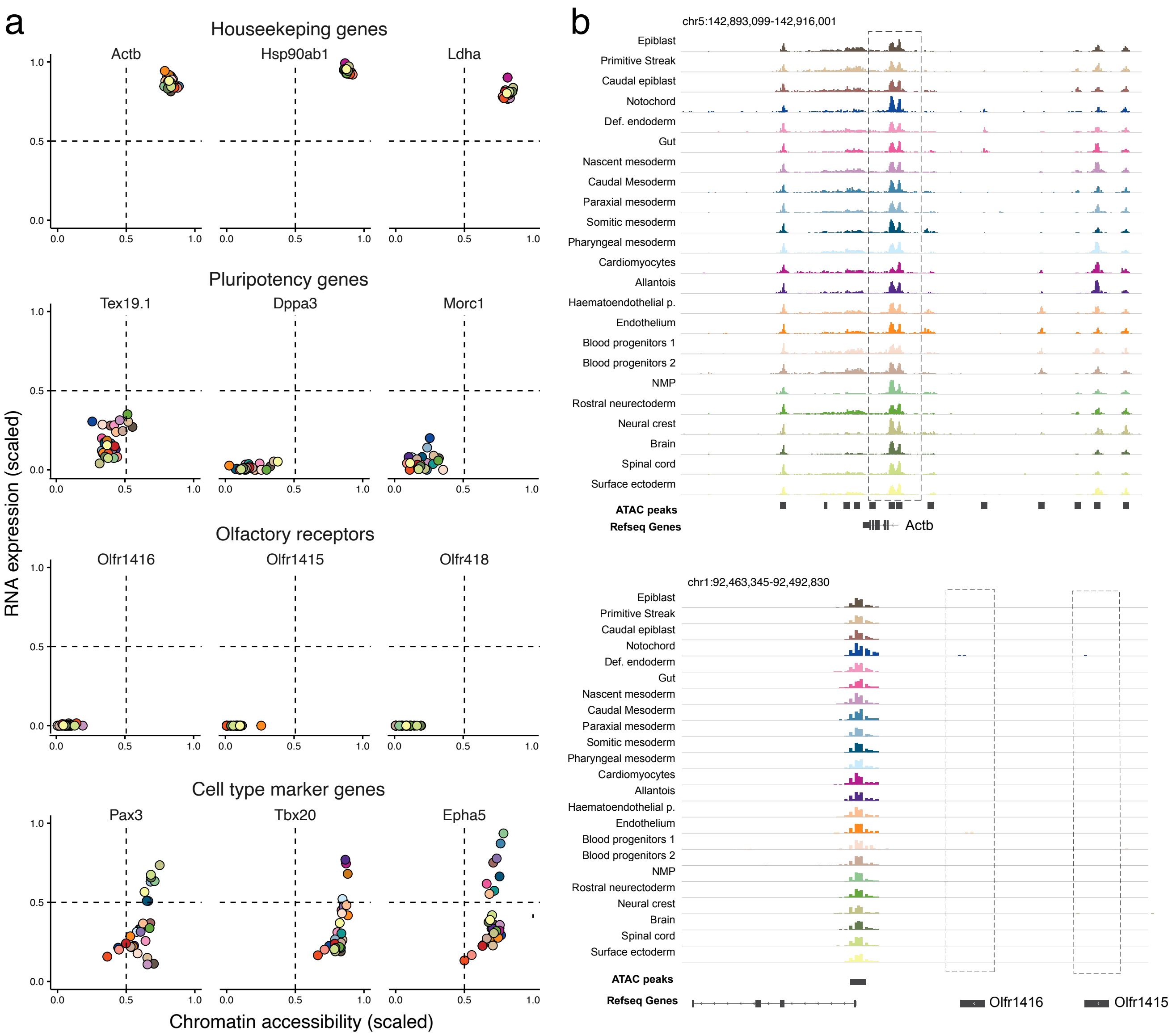


**Figure S6: Representative examples of RNA expression and chromatin accessibility values for different gene sets.**

(a) RNA expression and promoter chromatin accessibility values of different genes quantified for each cell type. The first row shows examples of housekeeping genes (positive control, highly expressed genes with open chromatin). The second row shows examples of naive pluripotency genes. The third row shows examples of olfactory receptors (negative control, non-expressed genes with closed chromatin). The fourth row shows examples of cell type marker genes.

(b) Genome browser snapshots displaying the *Actb* loci (housekeeping gene) and the *Olfr1416* loci (olfactory receptor). Each track displays pseudobulk ATAC-seq signal for a given celltype.

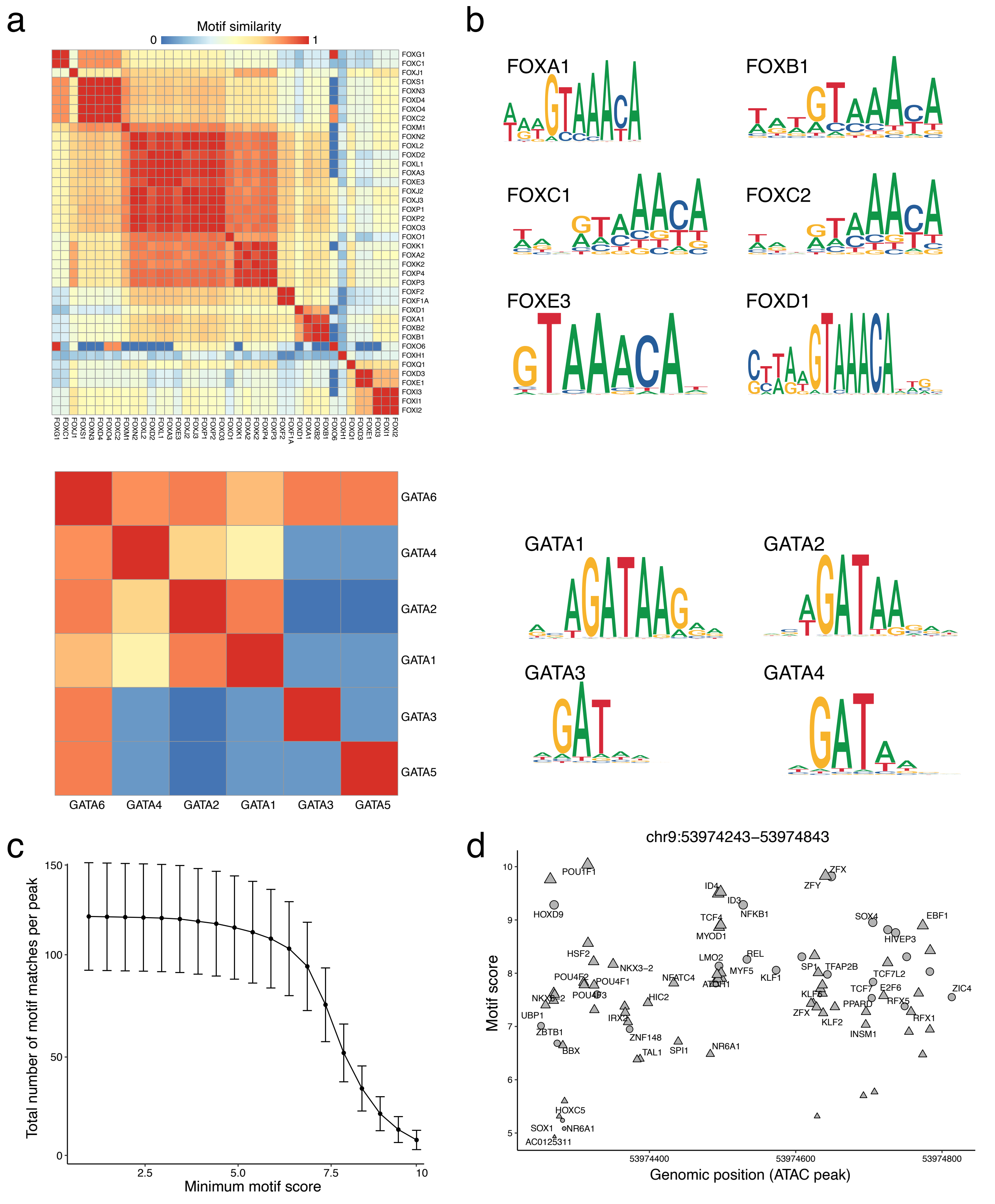

**Figure S7: Transcription Factor motif similarities poses challenges for studying gene regulation.**

(a) Heatmap showing the similarity between motif sequences for FOX (top) and GATA motifs (bottom). The similarity score is a normalised version of the sum of column correlations proposed in (Pietrokovski 1996). A score of 1 is expected for two identical motifs, whereas a score of 0 is expected for unrelated motifs.

(b) Representative examples of FOX (top) and GATA (bottom) transcription factor motifs to illustrate the similarity between motifs from the same TF family.

(c) Number of TF motifs per peak as a function of the minimum TF motif score cutoff.

(d) Location of motifs within a representative ATAC peak. Each dot represents a TF motif match within the genomic sequence. The x-axis displays the position of the match and the y-axis displays the TF motif score.

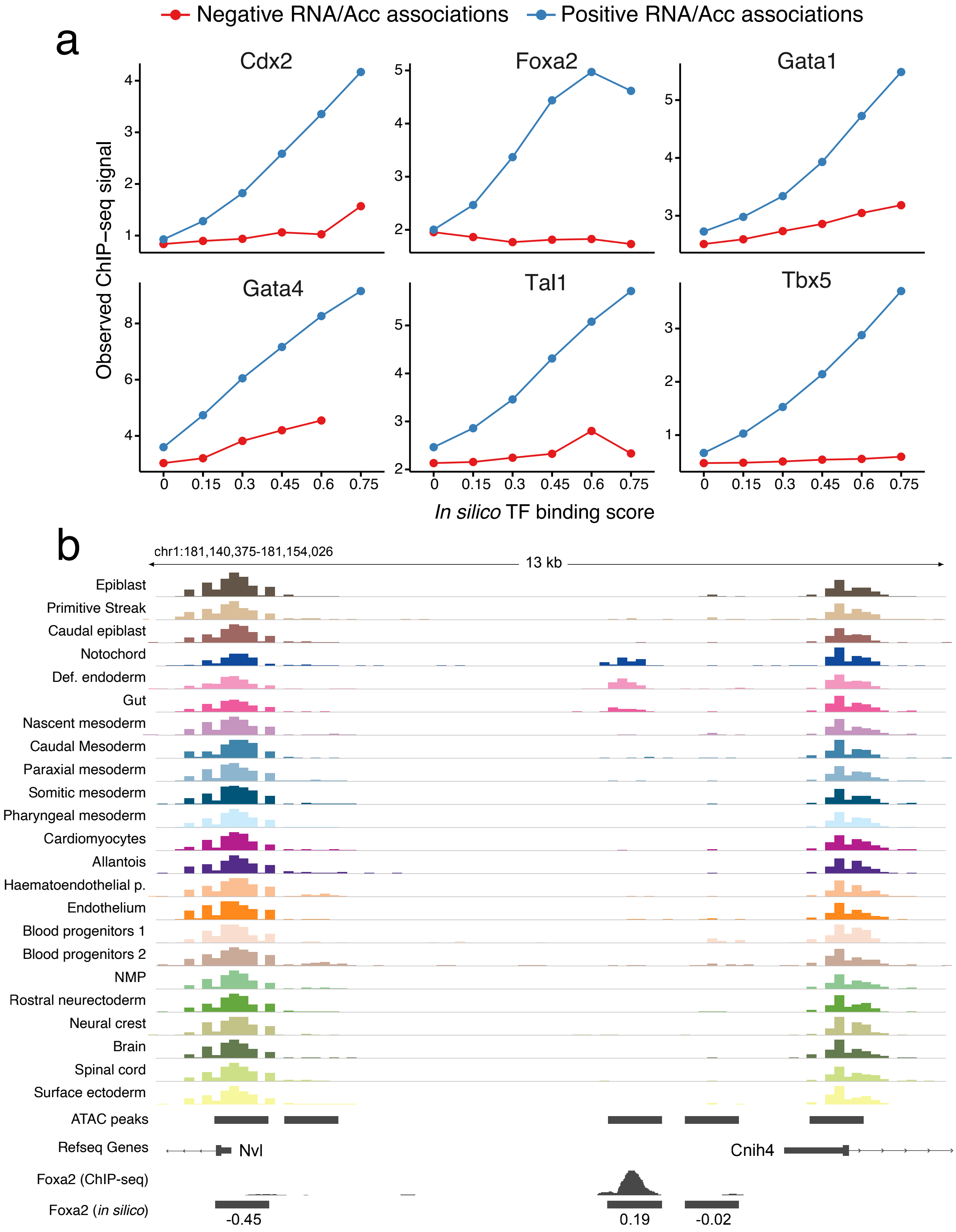


**Figure S8: Negative *in silico* TF binding scores that represent repressive interactions between TFs and chromatin accessibility changes display low consistency with ChIP-seq data.**

(a) Experimental ChIP-seq signal (y-axis) as a function of the *in silico* TF binding score (x-axis), split by the sign of the correlation between RNA expression and ATAC peak accessibility.

(b) Representative genome browser snapshot showing a locus with predicted Foxa2 binding sites. Each track displays pseudobulk ATAC-seq signal for a given cell type. The bottom tracks display ChIP-seq for Foxa2 binding (as a validation) and the *in silico* predicted binding sites for Foxa2.

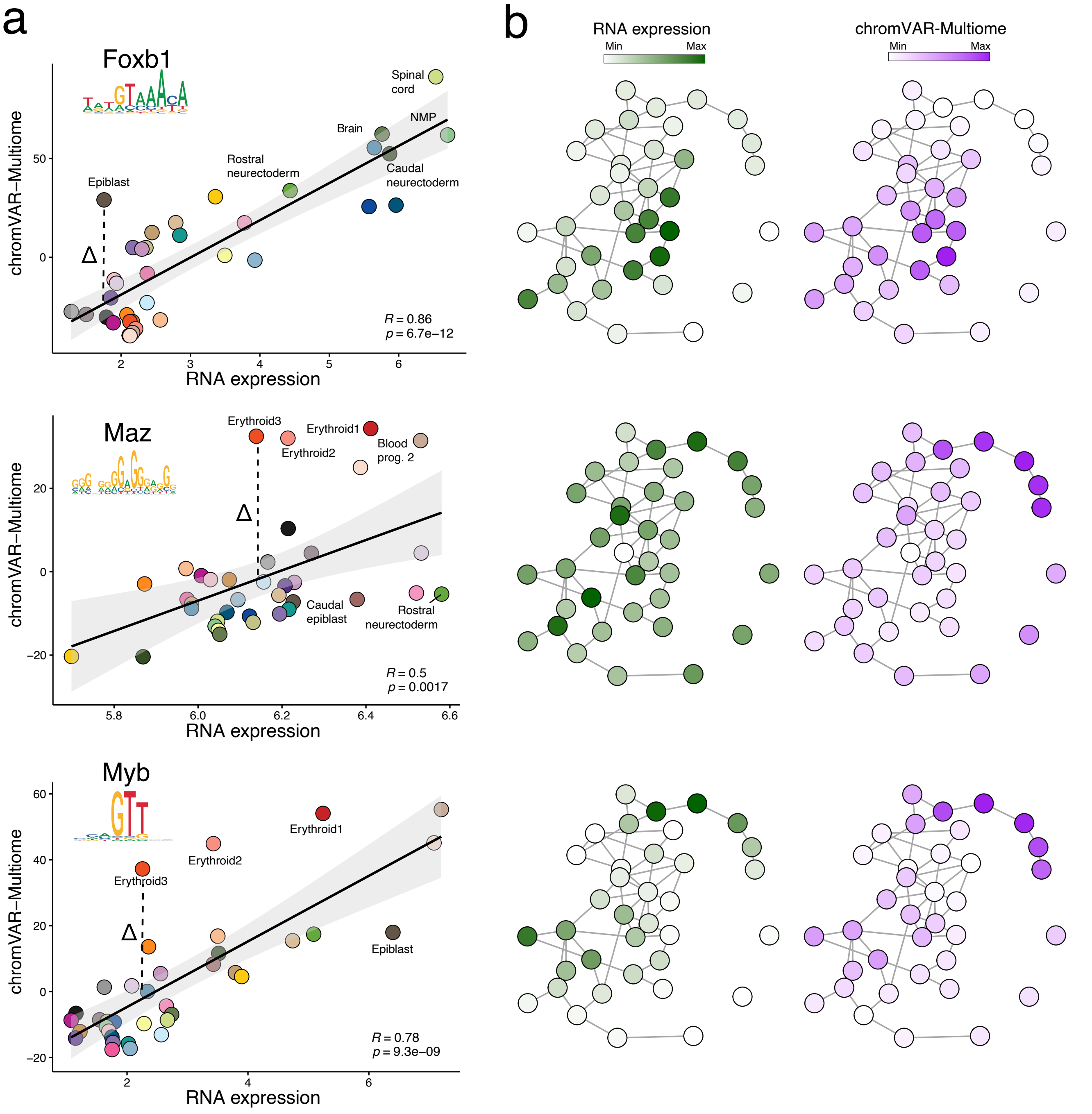


**Figure S9: Transcription factor chromatin activity scores infered with chromVAR-Multiome can be used to study the coordination between gene expression and chromatin accessibility dynamics.**

(a) Scatter plots show the correlation between the TF's RNA expression and the chromatin accessibility of target regions, quantified at the pseudobulk level using chromVAR-Multiome. Each dot corresponds to a different cell type. Highlighted are the residual values for selected cell types that result from a linear model linking gene expression and chromVAR-Multiome scores. High positive residuals represent events where the chromatin accessibility of TF targets for a cell type is larger than expected given the TF expression values, and hence can represent events where epigenetic priming might occur. Shown in the bottom right corner is the Pearson's correlation coefficient with associated p-value.

(b) PAGA representation of cell types (as in Figure 1b), where each node is coloured by the TF's expression (green) and the corresponding chromVAR-Multiome scores (purple).

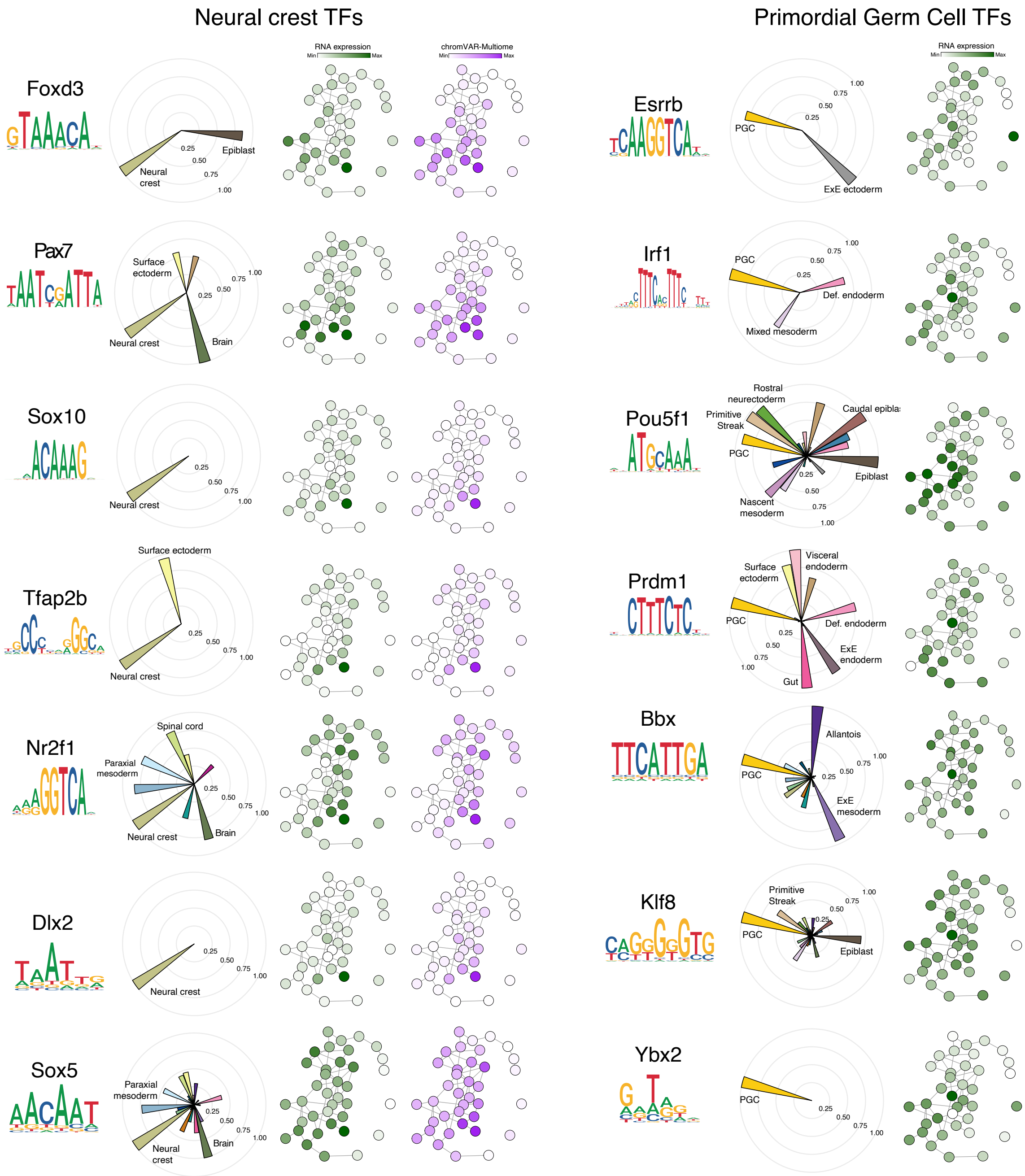


**Figure S10: Overview of Transcription Factor activities for Neural Crest and Primordial Germ Cells markers.** Each row shows the TF activities that result from performing differential analysis of the chromVAR-Multiome values (Methods). The higher the score for TF *i* in celltype *j*, the more active TF *i* is predicted to be in cell type *j*, with a minimum score of 0 and a maximum score of 1. Each panel shows: Transcription Factor (TF) of interest, alongside its DNA motif (left). Polar plots displaying the TF activity scores for each cell type (Middle). PAGA representation of cell types (as in Figure 1b) with each node coloured by the gene expression level (green) and chromVAR-Multiome score (purple) (Right). TF markers for Neural Crest are shown in the first column and TF markers for Primordial Germ cells (PGCs) are shown in the second column.

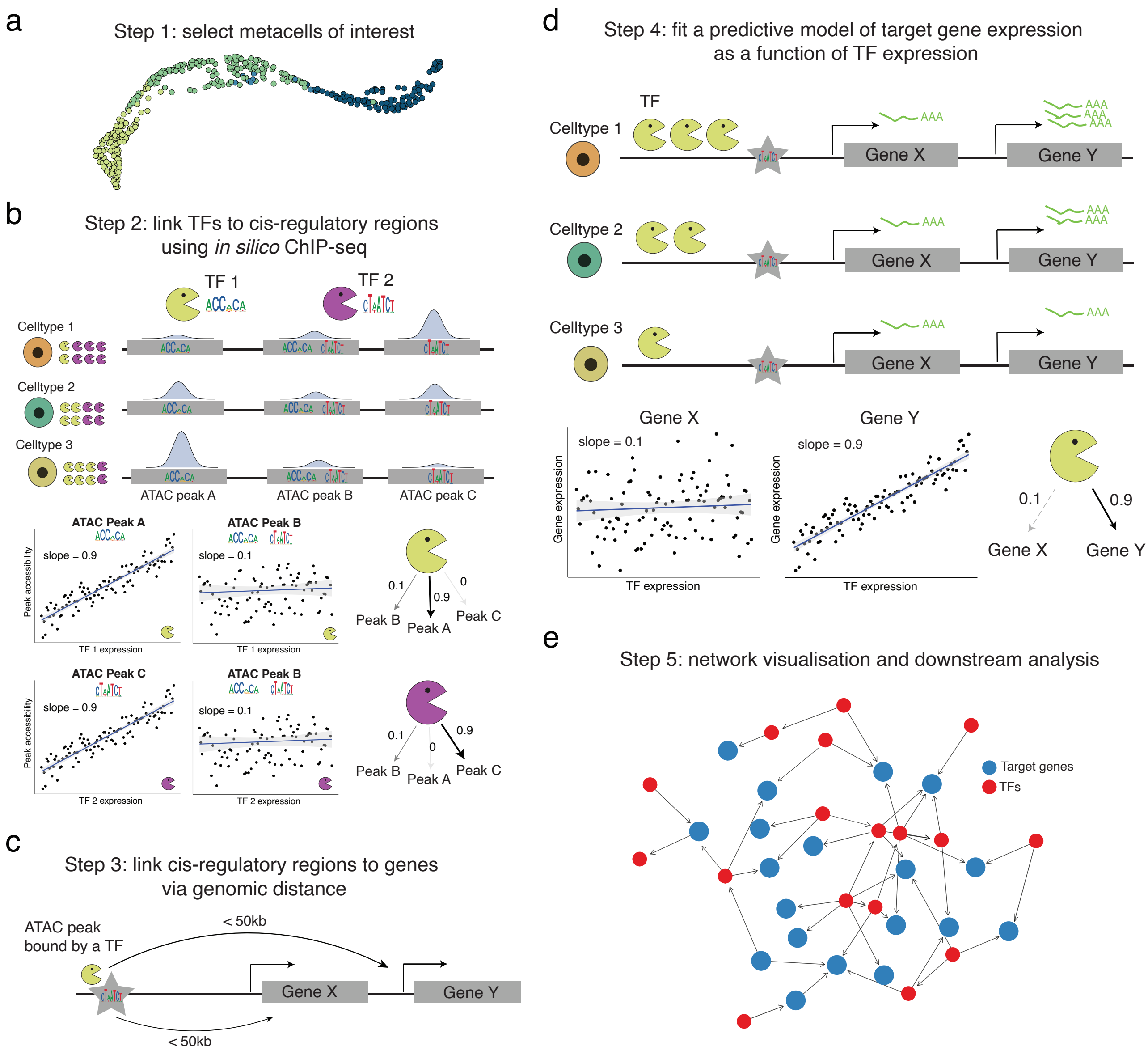


**Figure S11: Schematic of the methodology for Gene Regulatory Network (GRN) inference.**

(a) The first step is to select metacells of interest. We discourage the use of single cell resolution, as the sparsity of scATAC-seq makes it challenging to obtain reliable association estimates between the RNA expression of Transcription Factors (TFs, which are typically lowly expressed genes) and chromatin accessibility of target cis-regulatory elements. We refer the reader to the Supplementary Information for a more detailed discussion.

(b) The second step is to use the *in silico* ChIP-seq methodology to link TFs with cis-regulatory elements. This is the same diagram as shown in Figure 2a. Note that the *in silico* ChIP-seq results will vary depending on the metacells that are used as input.

(c) The third step is to link cis-regulatory regions that are predicted to be bound by TFs to nearby genes via genomic distance. Note that this is a many-to-many mapping, where each gene can be linked to multiple cis-regulatory regions, and each cis-regulatory region can be linked to many genes.

(d) The fourth step is to build a predictive model of target gene RNA expression as a function of the TF's RNA expression. Although some GRN inference methods have used non-linear regression models, here we use linear regression models, as they provide more stable, interpretable and generalisable estimates.

(e) The final step is to visualise the GRN as directed graph and perform quantitative analysis on the network. Red notes represent TFs, whereas blue nodes represent target genes. The edge width is given by the slope of the linear models.

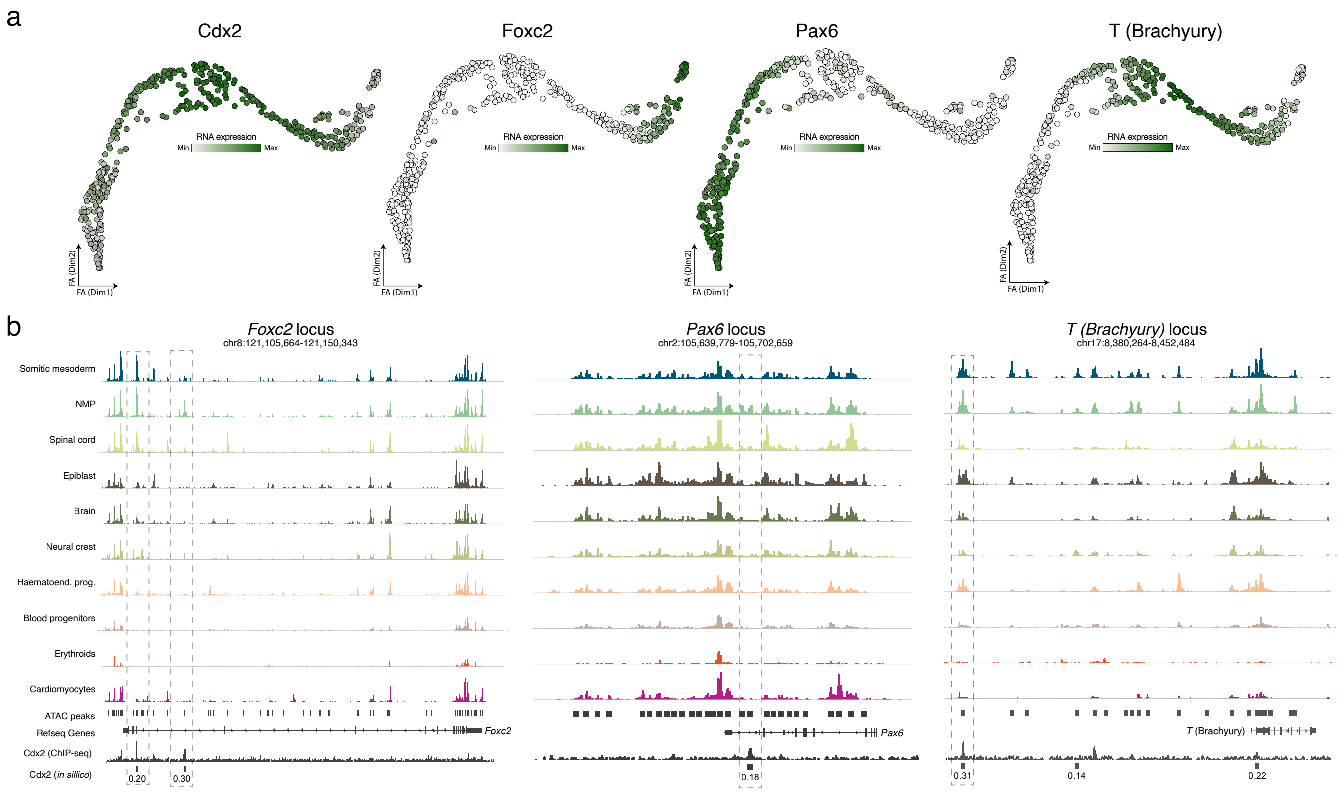


**Figure S12: Examples of repressive interactions between Cdx2 and other TFs that specify Spinal cord and Somitic mesoderm fate.**

(a) Force-atlas layout of the NMP differentiation trajectory. Each dot corresponds to a metacell, coloured by the RNA expression of Cdx2, Foxc2, Pax6 and Brachyury (T).

(b) Genome browser snapshot of different loci that code for genes associated with Somitic mesoderm fate (Foxc2 and T) and Spinal cord fate (Pax6). Each track displays pseudobulk ATAC-seq signal for a given celltype. Shown in the bottom is the *in silico* ChIP-seq predictions for Cdx2 and the experimental ChIP-seq signal for Cdx2 profiled in NMP-like cells (Amin et al. 2016). Highlighted are cis-regulatory elements that represent Cdx2 binding sites based on computational predictions and experimental data.

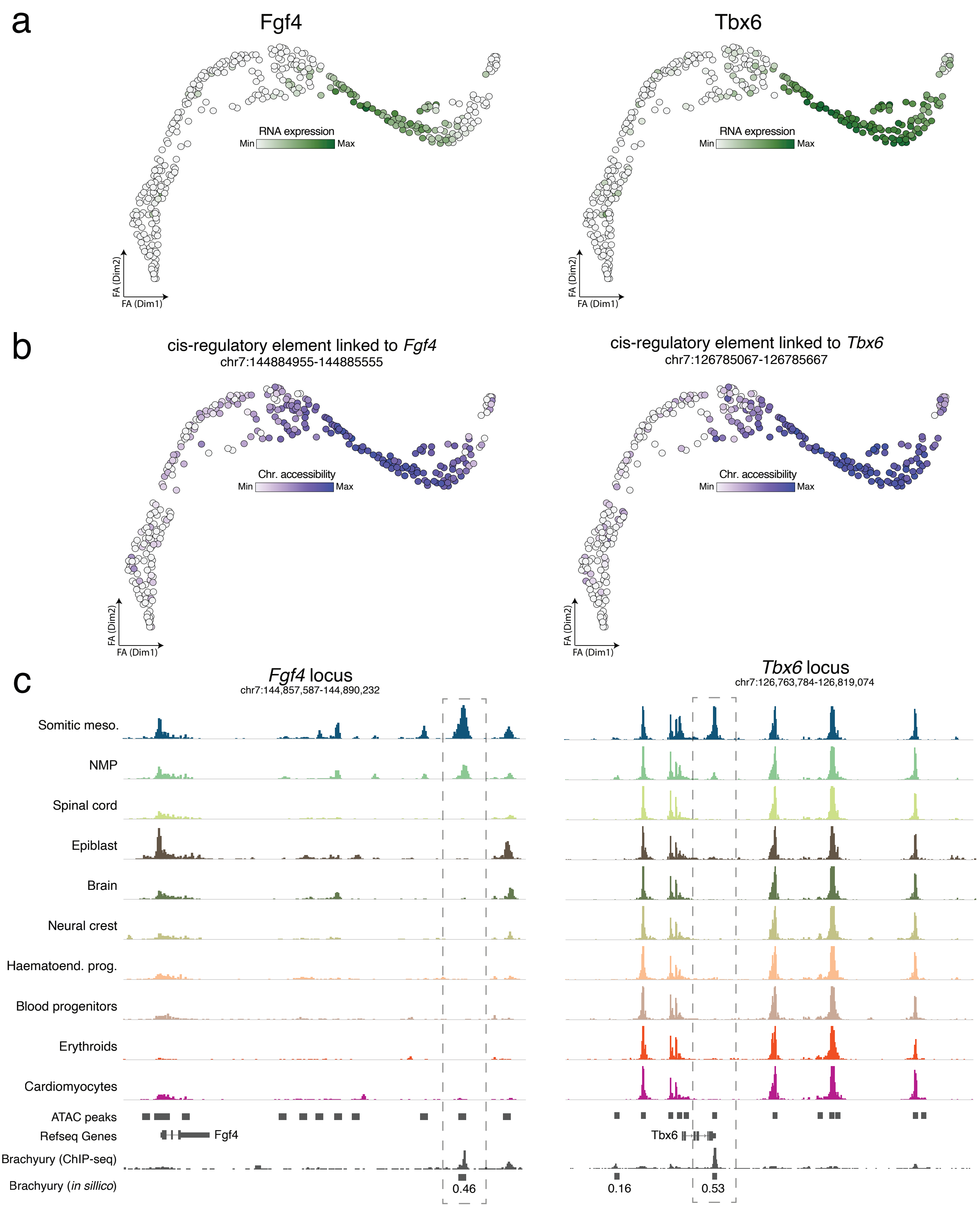


**Figure S13: Representative examples of cis-regulatory elements targeted by Brachyury that display epigenetic priming in NMP cells**

(a) Force-atlas layout of the NMP differentiation trajectory. Each dot corresponds to a metacell, coloured by the RNA expression of Fgf4 and Tbx6, respectively.

(b) Force-atlas layout of the NMP differentiation trajectory. Each dot corresponds to a metacell, coloured by the chromatin accessibility of cis-regulatory regions linked to *Fgf4* and *Tbx6*, respectively. Note that chromatin accessibility of the cis-regulatory regions precede the expression of the associated genes, shown in panel (a).

(c) Genome browser snapshot of chromatin accessibility signal around the loci that encode two genes linked to Somitic mesoderm fate: *Fgf4* and *Tbx6*. Each track displays pseudobulk ATAC-seq signal for a given celltype and genotype (WT in blue and Brachyury KO in red). Shown in the bottom are the *in silico* ChIP-seq predictions for Brachyury and the experimental ChIP-seq signal for Brachyury profiled in Embryoid Bodies (Tosic et al 2019). Highlighted are the Brachyury-targeted cis-regulatory elements shown in (b).

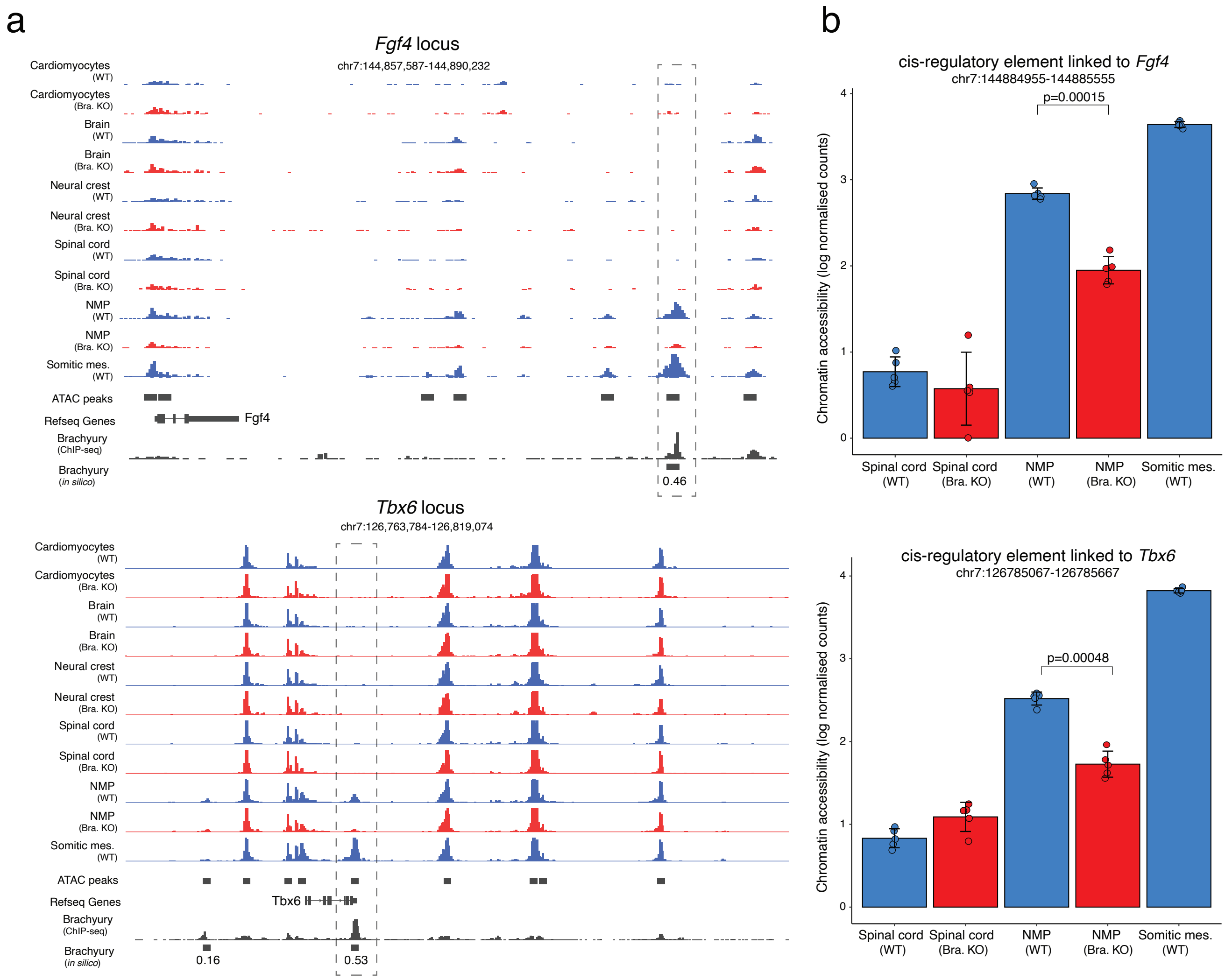


**Figure S14: Representative examples of cis-regulatory elements that display impaired epigenetic priming in Brachyury KO NMP cells.**

(a) Genome browser snapshot of chromatin accessibility signal around the loci that encode two genes linked to Somitic mesoderm fate: *Fgf4* and *Tbx6*. Each track displays pseudobulk ATAC-seq signal for a given celltype. Shown in the bottom are the *in silico* ChIP-seq predictions for Brachyury and the experimental ChIP-seq signal for Brachyury profiled in Embryoid Bodies (Tosic et al 2019). Highlighted are cis-regulatory elements targeted by Brachyury that are differentially accessible in WT and KO NMP cells. Note that these loci are the same shown in Figure S13.

(b) Bar plots display the chromatin accessibility of cis-regulatory regions per cell type and genotype, quantified at the pseudobulk level with replicates (Methods). Each dot corresponds to a pseudobulk replicate. Error bars display the standard deviation across replicates. Shown on top of the NMP bar plots is the p-value of a t-test comparing the mean accessibility between WT and KO NMP pseudobulk replicates.
