## Supplementary Table 1 for "Decoding gene regulation in the mouse embryo using single-cell multi-omics"

| **Supplementary Table 1. TF ChIP-seq experiments used to validate the *in silico* ChIP-seq** | | |
|---|---|---|
| **TF** | **GEO** | **Cell type** |
| CDX2 | GSE84899 | Cdx2 wtEpiSCs 24h |
| TAL1 | GSM1692848 | ES derived hemogenic endothelium |
| GATA1 | GSE69099 | ES derived hematopoietic progenitors |
| FOXA2 | GSE116258 | ES derived endodermal cells |
| GATA4 | GSE77548 | ES derived cardiomyocyte precursors |
| TBX5 | GSE77548 | ES derived cardiomyocyte precursors |
| RUNX1 | GSM1692856 | ES derived hematopoietic progenitors |
| NKX2-5 | GSM2054327 | ES derived cardiomyocyte precursors |
