## Supplementary Table 2 for "Decoding gene regulation in the mouse embryo using single-cell multi-omics"

### Supplementary Table 2. TFs previously linked to Neural crest and Primordial Germ Cell identity

| gene | celltype | evidence | species | references |
|---|---|---|---|---|
| DLX2 | Neural_crest | yes | zebrafish | https://doi.org/10.1016/j.ydbio.2007.11.005 |
| FOXD3 | Neural_crest | yes | mouse | https://doi.org/10.1242/dev.012179<br>https://doi.org/10.1242/dev.128.21.4127 |
| NR2F1 | Neural_crest | some | human | https://doi.org/10.1016/j.stem.2012.07.006 |
| SOX10 | Neural_crest | yes | mouse | https://doi.org/10.1016/s0012-1606(03)00161-1<br>https://doi.org/10.1038/ng0198-60 |
| SOX5 | Neural_crest | yes | chick | https://doi.org/10.1242/dev.01329 |
| TFAP2B | Neural_crest | yes | chick | https://dx.doi.org/10.1101%2Fgr.249680.119 |
| TFAP2C | Neural_crest | yes | zebrafish | https://doi.org/10.1016/j.ydbio.2006.12.042 |
| ETS1 | Neural_crest | yes | frog | https://doi.org/10.1074/jbc.m115.644864 |
| TFAP2A | Neural_crest | yes | chick | https://dx.doi.org/10.1101%2Fgr.249680.119 |
| PAX3 | Neural_crest | yes | frog | https://doi.org/10.1073/pnas.1219124110 |
| SOX9 | Neural_crest | yes | chick | https://doi.org/10.1242/dev.00808<br>https://doi.org/10.1242/dev.129.2.421 |
| MEF2C | Neural_crest | yes | mouse | https://doi.org/10.1016/j.devcel.2007.03.007 |
| PAX7 | Neural_crest | yes | mouse | https://doi.org/10.1371/journal.pone.0041089 |
| ALX1 | Neural_crest | yes | mouse | https://doi.org/10.3389/fcell.2022.777887 |
| ELK3 | Neural_crest | yes | chick | https://doi.org/10.1016/j.ydbio.2012.12.009 |
| TWIST1 | Neural_crest | yes | mouse | https://doi.org/10.1371/journal.pgen.1003405 |
| HBP1 | PGC | weak | mouse | https://doi.org/10.1002/dvdy.20053 |
| IRF1 | PGC | yes | mouse | https://doi.org/10.1038/s41467-021-27172-0 |
| ISL2 | PGC | no | NA | NA |
| PRDM1 | PGC | yes | mouse | https://doi.org/10.1038/nature03813<br>https://doi.org/10.1242/dev.01711<br>https://doi.org/10.1038/nature12417 |
| YBX2 | PGC | no | NA | NA |
| KLF8 | PGC | no | NA | NA |
| TFAP2C | PGC | yes | mouse | https://doi.org/10.1038/nature12417 |
| BBX | PGC | no | NA | NA |
| ESRRB | PGC | yes | mouse | https://doi.org/10.1016/j.mod.2004.01.006 |
| KLF2 | PGC | no | NA | NA |
| NR5A2 | PGC | yes | mouse | https://doi.org/10.1038/s41467-018-06230-0 |
| PGR | PGC | no | NA | NA |
| ID4 | PGC | no | NA | NA |
| KLF5 | PGC | no | NA | NA |
| NR3C1 | PGC | no | NA | NA |
| POU5F1 | PGC | yes | mouse | https://pubmed.ncbi.nlm.nih.gov/18395706/ |
| PRDM14 | PGC | yes | mouse | https://doi.org/10.1038/nature12417 |
